# Annotating metabolite mass spectra with domain-inspired chemical formula transformers

**DOI:** 10.1101/2022.12.30.522318

**Authors:** Samuel Goldman, Jeremy Wohlwend, Martin Stražar, Guy Haroush, Ramnik J. Xavier, Connor W. Coley

## Abstract

Metabolomic studies have succeeded in identifying small molecule metabolites that mediate cell signaling, competition, and disease pathology in part due to large-scale community efforts to measure mass spectra for thousands of metabolite standards. Nevertheless, the vast majority of spectra observed in clinical samples cannot be unambiguously matched to known structures, suggesting powerful opportunities for further discoveries in the dark metabolome. Deep learning approaches to small molecule structure elucidation have surprisingly failed to rival classical statistical methods, which we hypothesize is due to the lack of in-domain knowledge incorporated into current neural network architectures. We introduce a new neural network driven workflow for untargeted metabolomics, Metabolite Inference with Spectrum Transformers (MIST), to annotate mass spectrometry peaks with chemical structures generalizing beyond known standards. Unlike other neural approaches, MIST incorporates domain insights into its architecture by forcing the network to more directly link peaks to physical atom representations, neutral losses, and chemical substructures. MIST outperforms both standard neural architectures and the state-of-the-art kernel method on fingerprint prediction from spectra for over 70% of metabolite standards and retrieves over 66% of metabolites with equal or improved accuracy, with 29% strictly better. We further demonstrate the utility of MIST in a prospective setting to identify new differentially abundant metabolite structures from an inflammatory bowel disease patient cohort and subsequently annotate dipeptides and alkaloid compounds without spectral standards.

## 1 Main

Untargeted metabolomics is an important tool in advancing our understanding of cellular and environmental biochemistry [1, 2, 3, 4, 5, 6]. Liquid-chromatography tandem mass spectrometry (LC-MS/MS) is the primary experimental approach to conduct such analyses to identify new and often important metabolites: molecules are separated by column chromatography, ionized and measured based on mass (MS1), and fragmented by a higher energy collision (MS2) to produce a set of charged fragments with various mass-to-charge ratios, measured as a spectrum of peaks [7]. The analysis of these experiments is critically bottlenecked by an inability to accurately annotate these observed fragmentation spectra with the chemical structures of the molecules that produced them, with as many as 98% remaining unannotated [8]. In the many cases where the spectrum does not closely resemble a known standard spectrum, imperfect computational tools must be used to infer properties of the unknown molecule. Improving this single inference step has the potential to drastically increase the information gleaned across all routine untargeted metabolomics experiments [7].

Despite the recent breakthroughs in deep learning that have “neuralized” the adjacent protein structure prediction field [9, 10], the same cannot be said for metabolomics. Leading computational metabolomics annotation tools still rely upon hand-crafted heuristics and kernel functions (i.e., quantitative functions for comparing properties of mass spectra). Such methods can broadly be grouped into three categories: networking, forward prediction, and inverse prediction. Networking methods such as feature-based molecular networking [11, 12] cluster similar spectra to find neighborhoods of similar compounds (e.g., bile acids [13]). Forward methods (e.g., MetFrag [14, 15] and CFM-ID [16, 17]) augment the library of known metabolite standards by *in silico* fragmentation of molecules to generate artificial data. On the other hand, inverse fragmentation methods (e.g., CSI:FingerID [18] and FingerID [19]) predict molecular properties or the molecule structure from the spectra directly.

Inverse prediction of a molecule’s structural fingerprint from its spectrum is an entrenched paradigm in metabolomics, most recently forming the backbone of the winning solution at Critical Assessment of Small Molecule Identification 2022 (CASMI2022) [20, 21] (Fig. 1a). In particular, CSI:FingerID, the state of the art method assigns each peak in the fragmentation spectrum a chemical formula and arranging these in a fragmentation tree [22] (Fig. 1b-c). A set of kernels are then used to train a support vector machine to predict individual molecular fingerprint property bits for an input molecule. These predicted fingerprints, which often capture the presence or absence of particular molecular substructures, can then be queried against a database to annotate the spectrum with a specific molecular structure [18]. Subsequent models use these predicted fingerprints in various ways: CANOPUS predicts the chemical class of a target molecule from fingerprint [23], NPLinker associates a compound to its respective gene cluster in a genome [24], MSNovelist *de novo* generates a molecular structure from the putative fingerprint [25], COSMIC calibrates the uncertainty of molecular annotations [26], and Qemistree can be used to cluster compounds [27] (Fig. 1a). While great strides have been made in these downstream analyses, performance for all such tools is fundamentally limited by the first step of fingerprint prediction, which we aim to improve with this work.

**Fig. 1.**
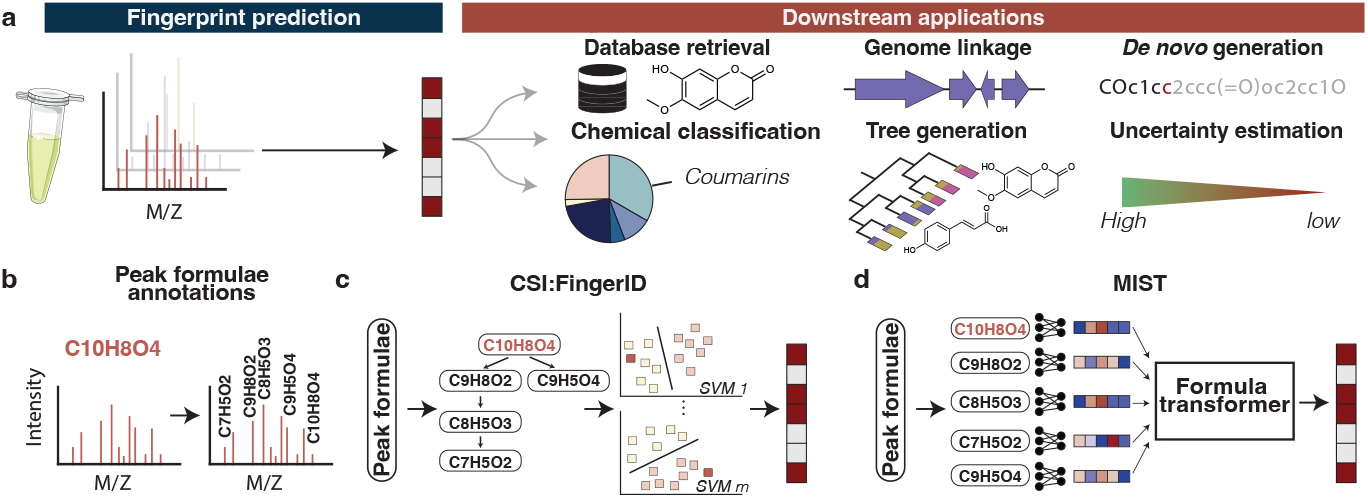
MIST replaces the critical spectrum-to-fingerprint prediction step in the computational workflow. **a.** A set of fragmentation peaks are extracted from a sample with tandem mass spectrometry. After predicting molecular property fingerprints, several different downstream software tools can be used to annotate molecules from databases, link metabolites to genomes, *de novo* generate plausible candidates, classify the unknown metabolite into compound classifications, generate statistics for all observed spectra, and calibrate certainty of annotations. **b.** The first step in the fingerprint prediction pipeline common to MIST and CSI:FingerID is to annotate the spectrum with a chemical formula for the full metabolite, then label each sub-peak with a chemical formula. **c.** CSI:FingerID arranges annotated sub-peak formulae into a “fragmentation tree”, and subsequently uses many independent support vector machine models to predict fingerprint property bits. **d.** MIST does not require a fragmentation tree, but rather directly utilizes chemical formulae as inputs to a formula transformer in order to predict molecular fingerprints.

Several efforts have been made to leverage deep learning to improve metabolomic analysis. Representation learning of spectra such as Spec2Vec [28], MS2DeepScore [29], and sinusoidal embeddings [30] can be used to learn a more meaningful distance between spectra to facilitate molecular networking. Forward models have incorporated feed forward networks and graph neural networks to predict a fragmentation spectrum directly from molecular structure [31, 32, 33]. In the inverse direction MSGenie [34], Spec2Mol [35], and MetFID [36] directly attempt to generate fingerprints or SMILES strings [37] from mass spectra, but no approach outperforms CSI:FingerID when trained with equivalent data. Deep kernel learning has also been leveraged to improve CSI:FingerID [38], but this approach still fundamentally relies upon decades of expert knowledge via hand-crafted input features and cannot be edited, retrained, or fine-tuned on new data in an independent study. We hypothesize that these neural network approaches are collectively unable to outperform their statistical counterparts because they lack in-domain knowledge in the architecture. By treating peak masses as discrete binned values, neural representations are less able to generalize between peaks and across examples.

Here, we introduce Metabolite Inference with Spectrum Transformers (MIST), a network based approach capable of outperforming current approaches despite not using hand crafted kernels. Rather than applying networks to binned spectra as prior neural models have, MIST first represents a spectrum as the set of chemical formulae of all peaks, borrowing inspiration from CSI:FingerID (Fig. 1d). We introduce inductive biases from the mass spectra domain: we implicitly featurize neutral loss relationships between fragments, simultaneously predict the structures of the metabolite and its fragments in each spectrum, use *in silico* forward augmentation to provide more training data for our model, and introduce a novel “unfolding” architecture to progressively increase the resolution of fingerprint predictions.

The ability of MIST to annotate spectra with structures from large virtual libraries of known biomolecules can be further enhanced through finetuned using contrastive representation learning. Excitingly, its learned latent embedding—a byproduct of the neural architecture—can also be used to cluster unannotated spectra with distances more representative of molecular distance than existing approaches. We show the utility of various model components and the inductive biases described above through thorough ablation studies on a publicly available dataset to provide a foundation for future model development. Finally, to demonstrate the prospective application of MIST to biological discovery, we analyze clinical samples from an inflammatory bowel disease patient cohort and propose new dipeptide and alkaloid structure annotations for differentially abundant metabolites.

MIST is distributed as an open source tool that can be easily integrated into existing pipelines, with or without retraining, and is freely available under the MIT license at ref. [39] and https://github.com/samgoldman97/mist.

## 2 Results

### 2.1 The MIST method

MIST accurately predicts fingerprints and compound annotations by using a deep neural network to learn a meaningful representation of an input mass spectrum. An input spectrum is composed of a single MS1 precursor mass and a list of mass over charge (m/z) peaks with corresponding intensity values. Rather than discretize the observed peaks into a fixed-length binned representation of intensities, MIST instead initially represents each spectrum peak as a chemical formula, which can be pre-labeled (inferred) based upon its mass-to-charge ratio using the SIRIUS algorithm [40] (Fig. 2a). MIST then directly transforms the representation at each peak by projecting these chemical formulae and intensities into feature vectors via a shallow feed forward multilayer perceptron (MLP). We posit that this “set of formulae” input representation is less noisy and includes more chemical intuition around likely atomic composition of each peak than binned spectra inputs used in other neural network models [36]. On the other hand, MIST does not utilize any information about the inferred relationship between peaks specified by the fragmentation tree from CSI:FingerID [18].

**Fig. 2.**
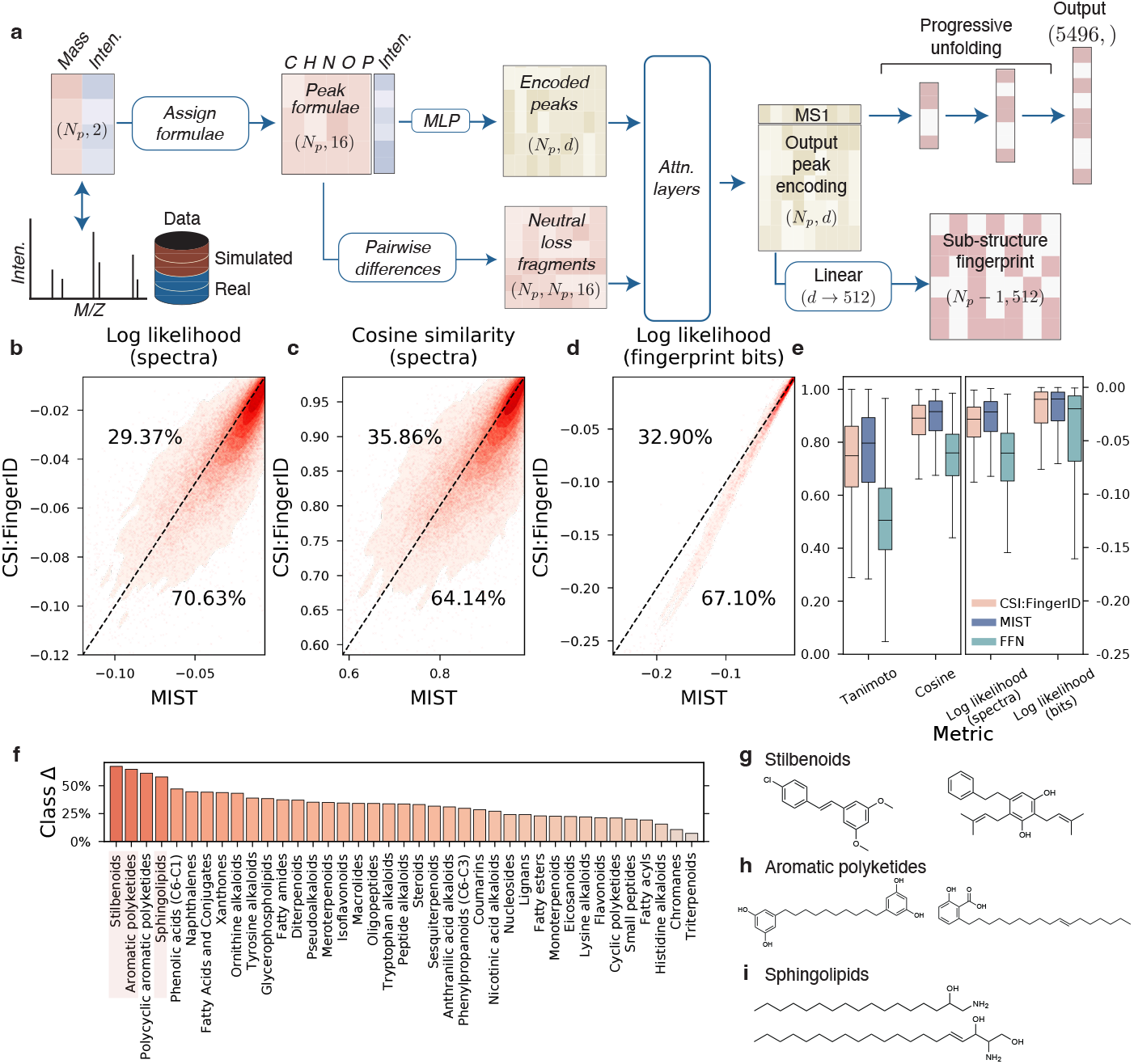
MIST accurately predicts compound fingerprints from mass spectra. **a.** Overview of MIST model architecture. An input spectrum of (*mass-to-charge, intensity*) pairs is transformed into chemical formulae vectors, encoded, and fed into a chemical formula transformer along with pairwise neutral losses between formulae. An unfolding module predicts full molecular fingerprints and a separate auxiliary module predicts substructure fingerprints as a secondary training signal. **b,c.** Molecular fingerprints are predicted by MIST and CSI:FingerID for every spectrum in the test set. The performance for each spectrum by cosine similarity to the true fingerprint (b) or log likelihood (c) is evaluated and plotted. Points below the line represent instances where MIST is more performant. **d.** Equivalent evaluation showing the likelihood of predicting each fingerprint bit correctly across all spectra. **e.** The performance of CSI:FingerID [18], MIST, and FFN, a baseline inspired by MetFID [19], are shown. “Tanimoto,” “Cosine,” and “Log likelihood (spectra)” indicate performance across spectra and “Log likelihood (bits)” indicates performance across bits (higher is better). Median lines are shown; boxes show the interquartile range, with fliers indicating 1.5x interquartile ranges. **f.** Performance differences by compound class as assigned by NPClassifier. All classes with > 40 molecules are shown. **g-i.** Example molecules from stilbenoid, aromatic polyketide, and sphingolipid classes. All results indicate predictions aggregated across 3 separate independent splits and model re-trainings. MIST fingerprints are created by averaging predictions from 5 separately trained models per split.

MIST learns relationships between multiple peaks and their inferred chemical formulae with successive Set Transformer multi-head attention layers [41, 42]. Given the importance of neutral loss fragments in expert analysis of spectra [43], we explicitly encode all pairwise neutral loss relationships as inputs to the attention layer modules. This allows MIST to update the initial representation at each of the peaks based solely on chemical formula with information about all other peaks and the pairwise losses between them. To obtain the final representation of the overall spectrum, we extract the learned representation of the peak corresponding to the precursor formula of the parent compound.

The spectral representation is intended to perform a variety of downstream tasks. The most essential is the prediction of the molecular fingerprint of the compound that generated the spectrum. Given the sparse nature of hashed fingerprints and how they are constructed, we introduce a novel “unfolding” layer where MIST progressively predicts the molecular fingerprint with larger and larger resolutions before reaching the final fingerprint prediction. This strategy enables the network to first learn coarse-grained molecular properties before generating the full resolution property vector, similar to how progressive growing has been used to generate high resolution images in computer vision [44].

Simultaneously, we ensure the spectral representation learns to make use of information reflected by each peak in the fragmentation pattern. A common strategy when training neural networks is to include auxiliary secondary tasks to increase performance. We use the MAGMa algorithm [45] to label chemical substructures for our training data and simultaneously train our model to predict fingerprints for substructures as an additional training signal to improve performance (Fig. S4)

A final key performance driver to MIST is the use of simulated data, inspired in part by the knowledge distillation strategy employed by AlphaFold [10, 46]. We train a forward neural network to predict spectra from molecules [31] and predict putative new spectra for a database of (unlabled) biological molecules (Fig. S3). These examples are randomly sampled in each epoch of model training to expand the dataset utilized for training. We refer the reader to Section 9.1 for a more complete description of the method.

### 2.2 Formula transformers outperform kernel baselines on fingerprint prediction

We directly compare MIST to the state of the art fingerprint prediction model, CSI:FingerID, within SIRIUS, to show the benefit of our approach. We prioritize direct comparison with CSI:FingerID as it was the highest performing method at CASMI2022 [20, 21], a prospective competition held in June 2022. MIST is trained on a dataset of positive ion spectra with hydrogen adducts constructed from NIST [47], MONA [48], and the GNPS databases [49], totaling 31,145 unique spectra of 27,797 unique compounds (Section 9.7). We train MIST to predict the 5496-dimensional fingerprint used by CSI:FingerID and evaluate performance on 3 separate splits of 20% data holdouts. We ensemble predictions from 5 separately trained MIST models with different random initializations; note that CSI:FingerID’s fingerprint prediction is itself an ensemble of 5496 separately trained models.

Of the 18,700 spectra in the test set holdouts, MIST predictions have higher cosine similarity to the true prediction for 11,994 of the examples (Fig. 2b). Further, despite training MIST to maximize cosine similarity of predicted fingerprints to true fingerprints rather than log likelihood directly, MIST fingerprint predictions still have higher log likelihoods for 64.12% of the spectra and 67.10% of the individual fingerprint bits (Fig. 2c,d). We observe similar trends when considering Tanimoto similarity of fingerprints when binarized at a 0.5 threshold (Fig. 2e). We also compare this to a more straightforward feed forward neural network (FFN) operating on binned (discretized) spectra, which performs substantially worse than both CSI:FingerID and MIST, indicating the importance of a more domain-inspired architecture (Fig. 2e, S2). Even with a single, non-ensembled model, MIST still performs better on over 55% of examples by cosine similarity (Fig. S5, Tab. S1). This performance advantage from MIST is consistent across chemical classes, with especially strong accuracy on stilbenoids, aromatic polyketides, and sphingolipids (Fig. 2f-i). Sphingolipids are especially important compounds in mediating host-microbe interactions, as bacteroides can synthesize sphingolipids with various carbon lengths and saturation patterns to alter host recognition. Novel modalities in sphingolipid metabolism will reveal more mechanisms of host-microbiome interactions [50].

### 2.3 Contrastive fine-tuning against decoy biomolecules allows for high accuracy metabolite annotation

Given the high accuracy of our fingerprint predictions, we turn our attention to the ability of MIST to successfully annotate the chemical structures of metabolites. In principle, one could directly query predicted fingerprints against a database of fingerprints for known biomolecules to retrieve the best matches. CSI:FingerID instead uses a series of custom Bayesian or heuristic distance functions to account for fingerprint bit correlations [51]. We focus on building a learned distance metric by fine-tuning MIST with a contrastive learning objective inspired by noise contrastive estimation [52] (Section 9.4) so that molecular fingerprints and spectra can be projected into a shared latent space; similar spectrum-compound pairs are trained to be close in terms of latent space distance (Fig. 3a). The contrastive MIST model is used to embed both a target spectrum and the molecular fingerprints corresponding to multiple potential metabolite annotations from a database such as PubChem.

**Fig. 3.**
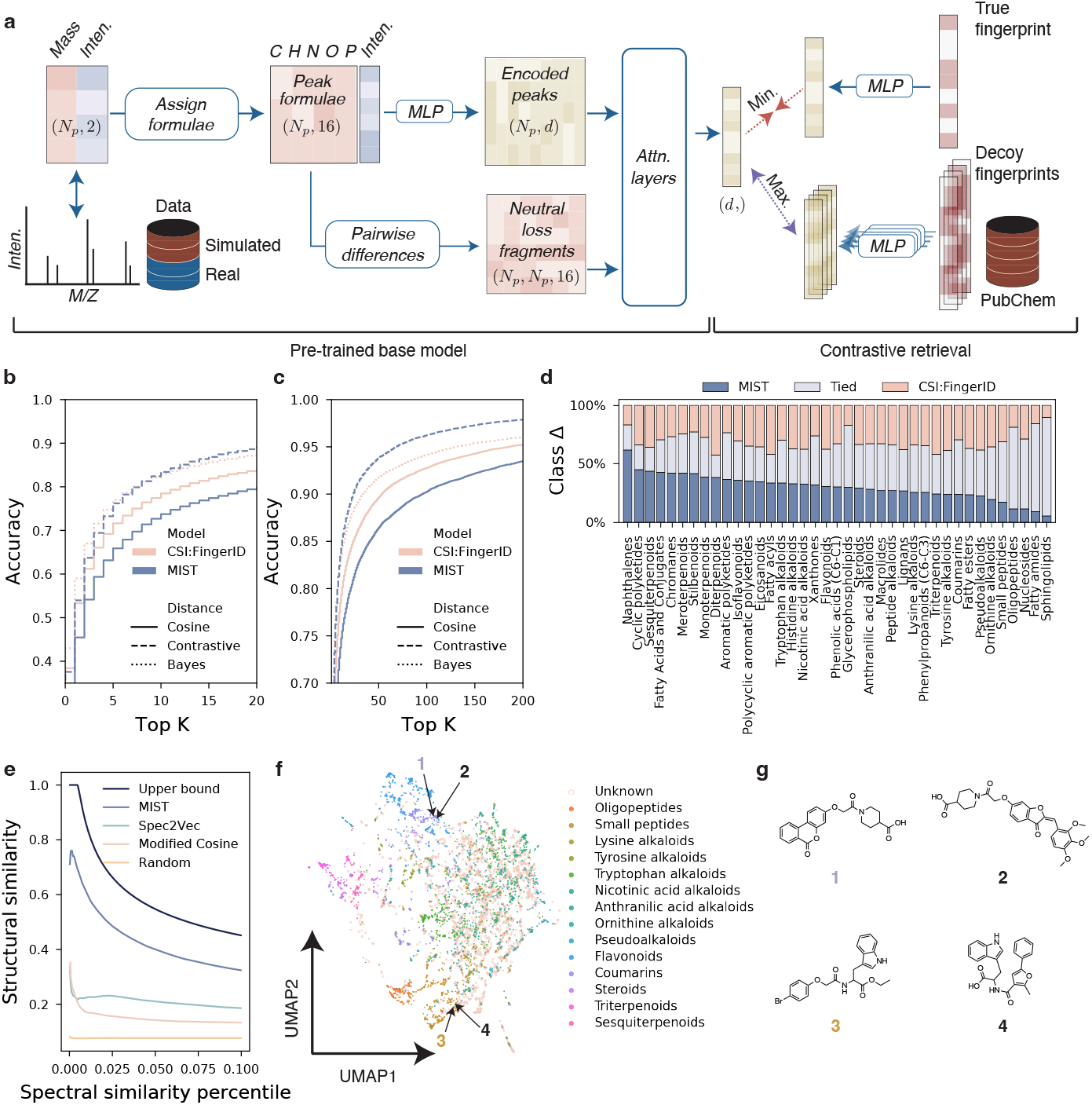
Contrastive fine-tuning improves compound annotation by database retrieval. **a.** Overview of the contrastive fine-tuning workflow for MIST. Model weights in the formula embedding module and attention layers are initialized from the fingerprint prediction tasks (“Pre-trained base model”) and fine-tuned such that fingerprint vectors are projected into the same latent hidden representation as the spectra are. The model minimizes distance between the spectra representations and their true structure’s representation and maximizes distance to decoy small molecules. **b,c.** MIST top-*k* retrieval accuracy assessed against CSI:FingerID by querying the PubChem reference database. Top-*k* accuracy is computed by ranking candidate compounds for each spectra and computing the fraction of spectra for which the true compound appears in the top-*k* rankings. Cosine distances use fingerprint predictions for retrieval, contrastive distances uses contrastive latent spaces, and “Bayes” is a custom CSI:FingerID fingerprint distance function. Accuracy is aggregated across 3 separate test folds of the data. **d.** Retrieval accuracy for all folds are segmented into various chemical classes. For each class for which > 40 examples exist, the fraction of examples on which MIST performs better, equivalently, or worse than CSI:FingerID are shown. **e.** MIST latent space spectra distances more effectively cluster high similarity compounds than Spec2Vec [28] distances or cosine distances. Test set compounds pairs are sorted by similarity according to various metrics and the average structural similarity of pairs is computed using Tanimoto distance at various percentiles for a single test fold. **f.** Contrastive spectra latent vectors for a single test fold are projected into 2D space with UMAP and colored by their respective compound class. Compounds 2 and 4 are example spectra for which NPClassifier fails to return an annotation. **g.** Example compounds from (f) reveal structural similarity of neighboring compounds in the UMAP space derived from MIST’s spectral embeddings.

MIST with contrastive embedding leads to a top-1 annotation accuracy of 37.39% on retrospective datasets, improving upon a 31.70% top 1 annotation accuracy if predicted fingerprints are used to retrieve candidate compounds directly (Fig. 3b; Tab. S2). Curiously, despite the improved fingerprint accuracy of our model in comparison to CSI:FingerID, we find that on the retrospective dataset, CSI:FingerID has equivalent top-1 accuracy (38.24%) compared to our method and higher top-1 accuracy when retrieving methods using a custom Bayesian fingerprint retrieval method [51]. It is possible that CSI:FingerID kernels, having been developed on this specific dataset, are overfit for this retrospective retrieval setting; prospective applications in previous CASMI competitions have exhibited lower accuracy [20]. A challenge for comparing such methods is that most annotation methods have been designed using nearly all public mass spectrometry data, so retrospective evaluation suffers from data leakage, requiring prospective studies to demonstrate utility. Because no handcrafted distance functions are utilized in our method and test set retrieval performance was not used to determine any parameters or hyperparameters, it is robust to this mode of bias.

We find that the contrastive distance function has especially strong performance at retrieving the correct compound within the top-*k* for *k* ≥ 20. For nearly all *k* ≥ 20, MIST retrieves the true structure more often than CSI:FingerID’s Bayesian retrieval (Fig. 3c, Tab. S2). This indicates that even when MIST predicts the top-1 molecule incorrectly, it still captures important details of the true compound.

We next investigate specific subclasses of chemical space for which MIST may have an advantage over CSI:FingerID. Overall, we find that MIST retrieves the compound on 66% of examples with equivalent or better rank than CSI:FingerID (strictly better on 29% of the data). Analyzing the fraction of examples at which MIST is better by chemical classes, we see that there is not a clear agreement with fingerprint prediction. Whereas MIST seems to have an advantage on fingerprint prediction within sphingolipids (Fig. 2f), we see that MIST and CSI:FingerID have near equivalent sphingolipid retrieval accuracy (Fig. 3d). We attribute this to the fact that certain chemical classes have higher top-1% accuracy, possibly due to the bias within the retrieval libraries themselves (e.g., 43/57 sphingolipids are retrieved correctly by both methods). Nevertheless, MIST offers clear improvements on important chemical classes such as sesquiterpenoids, which have long been known to have important therapeutic properties [53].

### 2.4 Neural embeddings elucidate compound class clusters

Given the increased retrieval accuracy at higher values of *k*, we next asked if MIST is capable of meaningfully organizing chemical space in its learned latent space representation of mass spectra. Molecular networking has long been an important tool to cluster spectra that may represent similar molecules, used recently to guide the discovery of new bile-acid conjugate molecules [12, 13]. A challenge of molecular networking is the choice of a meaningful spectral distance and distance cut-off to define network edges.

MIST learns contrastive embeddings with inter-spectra distances that highly correlate with metabolite structure similarity, establishing a new way to form molecular networks. We compare this effect directly against modified cosine distance as implemented in MatchMS [54] and Spec2Vec [28], a recent representation learning strategy for embedding spectra (Fig. 3e). For 38,850,289 pairs of spectra scored by spectral similarity, the average structural similarity of the top 0.1% according to MIST is 0.32 compared to a theoretical max of 0.45 using the top 0.1% of most similar pairs, whereas using Spec2Vec yields a lower average structural similarity of 0.19. By this measure, MIST’s spectral similarity metric represents a > 50% improvement in structural similarity from the current state of the art.

Rather than form molecular networks by drawing arbitrary distance cutoffs, the continuous nature of MIST’s embeddings also makes it amenable to dimensionality reduction via UMAP. Excitingly, we find that chemical classes of molecules as annotated by NPClassifier [55] are highly clustered in the UMAP space despite the model having no explicit representation of the molecular structure or its class when forming the UMAP (Fig. 3f). Further, because NPClassifier is itself a predictive algorithm, certain molecules remain unannotated. By manual inspection, we observe that even these unlabeled molecules are indeed quite structurally similar to their neighbors (Fig. 3g); while the unlabeled molecule **2** does not share the characteristic coumarin motif with molecule **1**, both molecules share a 4-piperidinecarboxylic acid substructure not detected by chemical classifications. Similarly, unlabeled molecule **4** bears a strong resemblance to its labeled peptide neighbor, **3** as both appear to be tryptophan derivatives.

### 2.5 MIST uncovers new dipeptides in clinical cohort data

To show the utility of MIST on real world data, we applied MIST to identify unknown metabolites from a recent clinical metabolomics dataset extracted from patients with ulcerative colitis (UC) and Crohn’s disease (CD) as reported by Mills et al. [56]. UC is an inflammatory bowel disease (IBD) characterized by continuous mucosal inflammation from the rectum to the proximal colon with an annual disease prevalence of up to 245 cases per 100,000 persons [57]. Mounting evidence has shown that the microbiome mediates UC treatment response and disease progression, including by the production of novel metabolites [58, 59, 60]. CD is similarly the second primary cause of IBD, presenting as a chronic and recurring inflammation with 214 cases per 100,000 persons in the United States [61].

Repeating the authors’ data processing pipeline, we extracted relative abundances for a total of 1,990 tandem mass spectra across the 210-patient clinical cohort (73 UC, 117 CD, 20 healthy), compared to an original 1,928 spectra extracted by Mills et al. We assigned chemical formulae to 1,788 spectra using the SIRIUS algorithm [40]. MIST annotated chemical structures for 1,499 of these spectra by searching in the Human Metabolome Database (HMDB) when possible and falling back to a larger search in PubChem for chemical formulae without candidate structures in HMDB. The use of Pub-Chem as a larger reference database added 1, 007 total spectra and metabolites to our analysis compared to the original study.

Following the original study, we sought to identify metabolomic correlates of disease severity as assigned by clinicians. Putative metabolites were grouped into chemical class using NPClassifier [55] and relative abundances for each chemical class were calculated for each patient. We identified correlation coefficients between the total abundance of each metabolite class and patient disease severity within the UC and CD cohorts. Dipeptides exhibited the highest correlation with disease severity for UC patients of any chemical class (*R* = 0.4776, *p*_adjust_ = 0.01) (Fig. 4a,b). This is consistent with class-level metabolite analysis from Mills et al. and their finding that increased activity of microbial proteases in dysbiosis led to dipeptide accumulation [56]. Peptides and amino acids are carbon sources for numerous inflammation-associated microbes, particularly oral cavity residents who have been observed to colonize and thrive in inflamed gut [62]. An understanding of such peptide structures could reveal nutrient niches and footprints of prevalent microbial proteases, possibly suggesting new disease biomarkers or therapeutic targets [63].

**Fig. 4.**
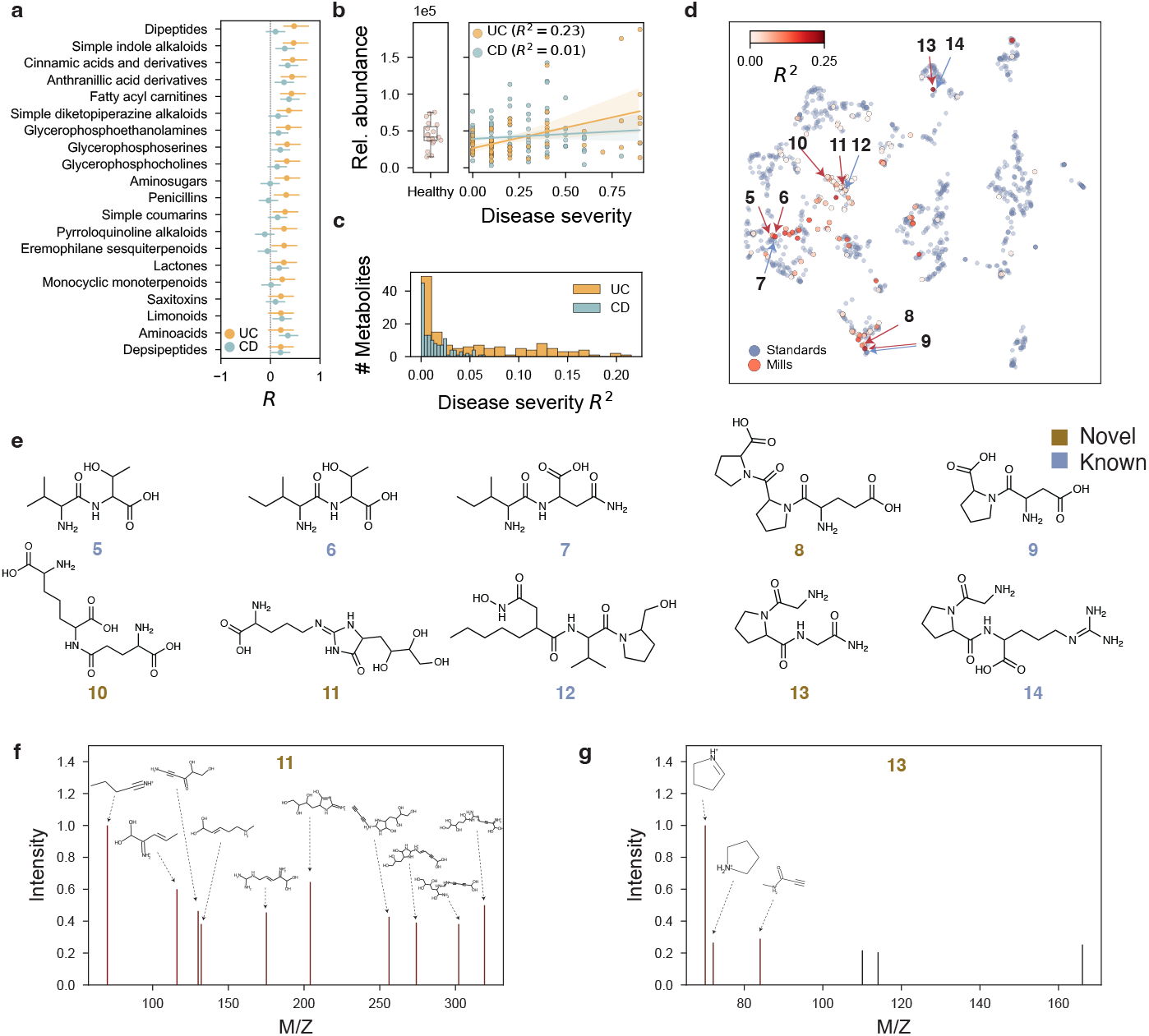
MIST discovers new putative and clinically relevant dipeptides. **a.** Metabolite classes ranked by their Pearson correlation with IBD disease severity. All putative metabolites are assigned disease classes with NPClassifier [55] and relative abundances are summed across classes. Within the ulcerative colitis (UC) and Crohn’s disease (CD) cohorts, metabolite classes are correlated with disease severity and sorted according to their correlation with UC (top 20 shown). **b.** Dipeptide relative abundances for each patient are plotted against disease severity within the healthy, UC, and CD cohorts. Dipeptides correlate with disease severity for UC patients (*R*^2^ = 0.23, *p* = 0.01) as observed by Mills et al. [56]. **c.** Within the Dipeptide class, the distribution of individual metabolite correlations are shown for both UC and CD cohorts. **d.** All putative dipeptide metabolites (red) are embedded into the latent space of a single, contrastive MIST model alongside embeddings for reference standard spectra from the original model training set and also labeled as dipeptides (blue). All embeddings are projected in 2D space with UMAP and plotted. Metabolites are colored by their coefficient of determination *R*^2^ with UC disease severity as computed in **c**. **e.** Annotations for example metabolites from **d** are shown across different compound clusters. Metabolites with chemical formulae not in standards library are colored brown. **f-g.** Example spectra and their MIST annotated compound are shown. Explained subpeaks are annotated using CFM-ID [16] and shown as validation of plausibility. Compound annotations are made using an ensemble of 5 MIST models, contrastive distance retrieval, and the HMDB reference database of molecules.

To interrogate these dipeptides, we next identified individual putative metabolite structures within the dipeptide class that most correlated with disease. At the individual metabolite level, most metabolites exhibited low correlation, whereas a smaller number had higher correlation with disease severity (Fig. 4c). To understand structural patterns driving these interactions, we embedded the spectra for known dipeptide standards alongside putative cohort dipeptide metabolites using a single MIST model and colored by correlation coefficient (Fig. 4d). Curiously, dipeptide metabolites tightly cluster within latent space into different groups; at least four such clusters (compounds highlighted with arrows) contain putative dipeptides that correlate with disease activity in the cohort. The cluster containing compounds **5**,**6**,**7** is well-characterized with little structural novelty, primarily composed of dipeptides with a single hydrophobic amino acid (e.g., valine, isoleucine) conjugated to a polar species (e.g., threonine) (Fig. 4e). On the other hand, the cluster containing compounds **8** and **9** showed proline conjugated to polar amino acids, with a pro-pro-glu tripeptide (**8**) not featured in the standards set demonstrating a high correlation. Because the chemical class annotation tool is itself a predictive model, certain tripeptides were observed, despite analyzing metabolites labeled as dipeptides. Interestingly, the cluster including **10**, **11**, and **12** shows high structural novelty, i.e., dissimilarity to known standards, but low correlation with disease severity.

We further inspected two of the novel spectrum annotations to determine whether MIST’s assigned structure can explain the observed spectrum using the CFM-ID [16] *in silico* fragmentation tool (Fig. 4f,g). For compound **11**, the novel putative species explains all of the top 10 peaks shown in the figure indicating a strong match (Fig. 4f). On the other hand, compound annotation **13** only explains half of the peaks observed in the spectrum (Fig. 4g). The three attributed peaks are due to the proline substructures, consistent with the nearby spectrum **14** annotation also containing a proline motif. Given that only three peaks of the six in the spectrum are explained, this example indicates that this may be a near match, rather than exact annotation, highlighting a potential shortcoming of the model and the importance of manual inspection.

While dipeptide metabolites correlated with disease severity within the CD and UC populations, they showed low differential abundance when compared to the healthy population itself (Fig. 4b). Few metabolite classes were differentially more abundant in the UC cohort compared to disease (Fig. 5a). Instead, we find that more metabolite classes are differentially *less* abundant in both UC and CD cohorts compared to healthy, including two classes that contain heterocyclic nitrogenous rings: piperidine (UC *p*_adjust_ = 0.034, CD *p*_adjust_ = 0.014) and pyridine (UC *p*_adjust_ = 0.006, CD *p*_adjust_ = 0.014) alkaloid classes (Fig. 5a-c). Novel immunoregulatory metabolites and their producers are continuously being uncovered, such as indoleacrylic acid, an inflammation-suppressing metabolite produced by commensal Peptostreptococcus [64]. This differential abundance indicates that these metabolite classes may harbor similar regulatory properties.

**Fig. 5.**
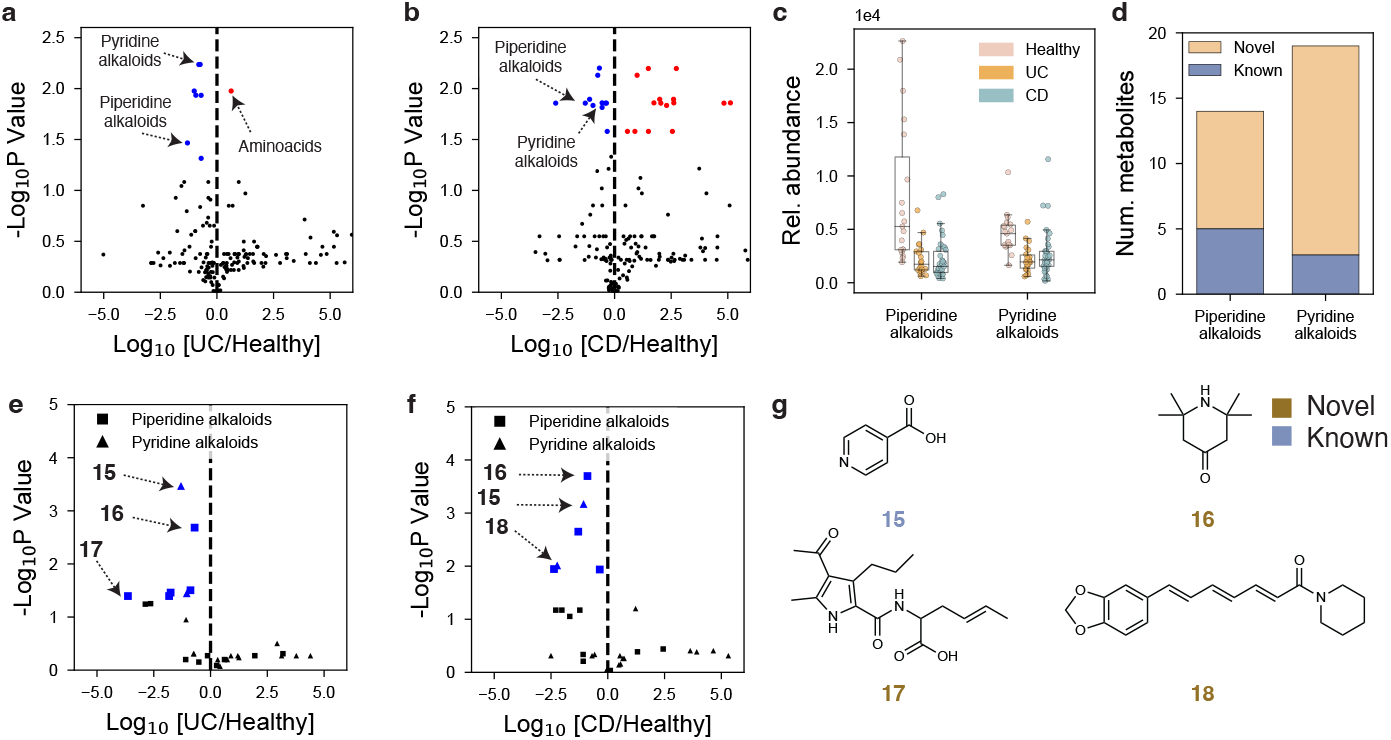
Putative alkaloids separate healthy and diseased IBD cohorts. **a,b.** Putative metabolite classes from the Mills et al. cohort [56] are scattered showing their fold change and respective p value comparing and ulcerative colitis (UC) (**a**) and Crohn’s disease (CD) (**b**) cohorts to the healthy cohort. **c.** Piperidine and pyridine alkaloids were selected as both are significantly less abundant in UC and CD cohorts. Distributions of patients’ relative abundances for these classes are shown for healthy, UC, and CD groups. **d.** The total number of putative metabolites in both significant classes are shown, highlighting the number of metabolites with putative novel structures not observed in the standards library. **e,f.** Individual metabolite abundance fold changes and statistical significance are shown comparing the healthy cohort to UC (**e**) and CD cohort (**f**). **g.** Select putative annotated molecule structures are shown for compounds **15** (isonicotinic acid, HMDB:0060665),**16** (triacetonamine, HMDB:0031779),**17** (unknown), and **18** (piperettine, HMDB:0034371). Spectra **15** and **16** are differentially less abundant for both UC and CD. Novel compounds indicate those without spectra standards and are labeled brown; compounds with respective standards are shown in blue. Compound annotations are made using an ensemble of 5 MIST models, contrastive distance retrieval, and the HMDB reference database of molecules. The top chemical classs annotation is found by running the top putative prediction through NPClassifier [55]. P values were computed using independent two-sided t-tests and adjusted for multiple hypothesis testing with the Benjamini Hochberg method. To have equal cohort sizes for healthy and diseased patients, UC and CD cohorts were subsetted to patients with disease severity score > 0.2.

Within the piperidine and pyridine alkaloid metabolite classes, we observed a total of 14 and 19 metabolites respectively, the majority of which were annotated as chemically novel structures by MIST (Fig. 5d). To better understand which metabolites drive the class-level effect, we analyzed differentially abundant metabolites within both alkaloid classes. We identified compounds **15**, **16**, **17**, and **18** as chemically diverse structures with significantly lower abundance than in healthy patients, with compounds **15** and **16** differentially less abundant across both disease cohorts (Fig. 5e-g). Curiously, not all compounds have the characteristic saturated and unsaturated six-member heterocycles expected for these alkaloid classes; compound **17** instead features a pyrrole ring. This again highlights that compound class annotations are predictive, even when the structure is known, further motivating manual inspection and the need to annotate compounds at the structure level, not class level.

Our results showcase how MIST can be used to identify and prioritize potential disease targets. Certain plant alkaloid compounds have been previously reported as ameliorating inflammation and could pose potential targets for future investigation [65]. Alternatively, compound **18**, piperettine (HMDB:0034371), is found naturally in peppers and certain spices. This suggests it may be a biomarker for dietary preferences patients follow in response to inflammation. Compound **15**, isonicotinic acid, is of similar interest as it has high structural similarity with niocotinic acid, a key intermediate in nicotine biosynthesis. Nicotine has a well documented and unknown protective effect in UC [66], indicating potential function for metabolite **15**. Future work may consider causal directions of these differential abundances.

### 2.6 Public data benchmarking as a community standard for future development

A critical impediment to the development of new models for structural elucidation in metabolomics has been the lack of community standards for benchmarking algorithms. While competitions like CASMI [67] provide test set data, they do not specify training datasets or level the playing field of access. In addition to comparing MIST directly to CSI:FingerID on the proprietary NIST [47] dataset and with proprietary fingerprints, we conduct a fully reproducible evaluation and ablation study of MIST on a 8, 030 spectra (7,131 unique compounds) subset of the GNPS dataset as prepared by Duhrkop *et al*. [23, 49].

Using this smaller but public dataset, we compare MIST first on the task of fingerprint prediction to variants of MIST that do not include pairwise neutral loss featurization (“MIST - pairwise”), use no auxiliary substructure losses (“MIST - MAGMa”), do not train on simulated fingerprints (“MIST - simulated”) or use only a feed forward neural network (“FFN”). We make predictions for 4096-bit circular Morgan fingerprints, which are straightforward to generate using RDKit [68]. As with the larger proprietary dataset, MIST sharply outperforms the FFN model in terms of cosine similarity and log likelihood (Fig. 6a,b; Tab. S3). Interestingly, we find that outside of using the overall chemical formula transformer, fingerprint unfolding has the largest effect on the performance of cosine similarity. Unlike in the case of CSI:FingerID fingerprint bit predictions, pairwise embeddings do not appear to improve performance of MIST, which may be a function of the limited public benchmarking dataset size. In support of this, we find that cosine similarity seems to be improving linearly as a function of dataset size in this regime (Fig. 6c). Log likelihood based evaluations showcase similar but noisier trends, as expected because the model is not explicitly trained to maximize log likelihood (Fig. 6d).

**Fig. 6.**
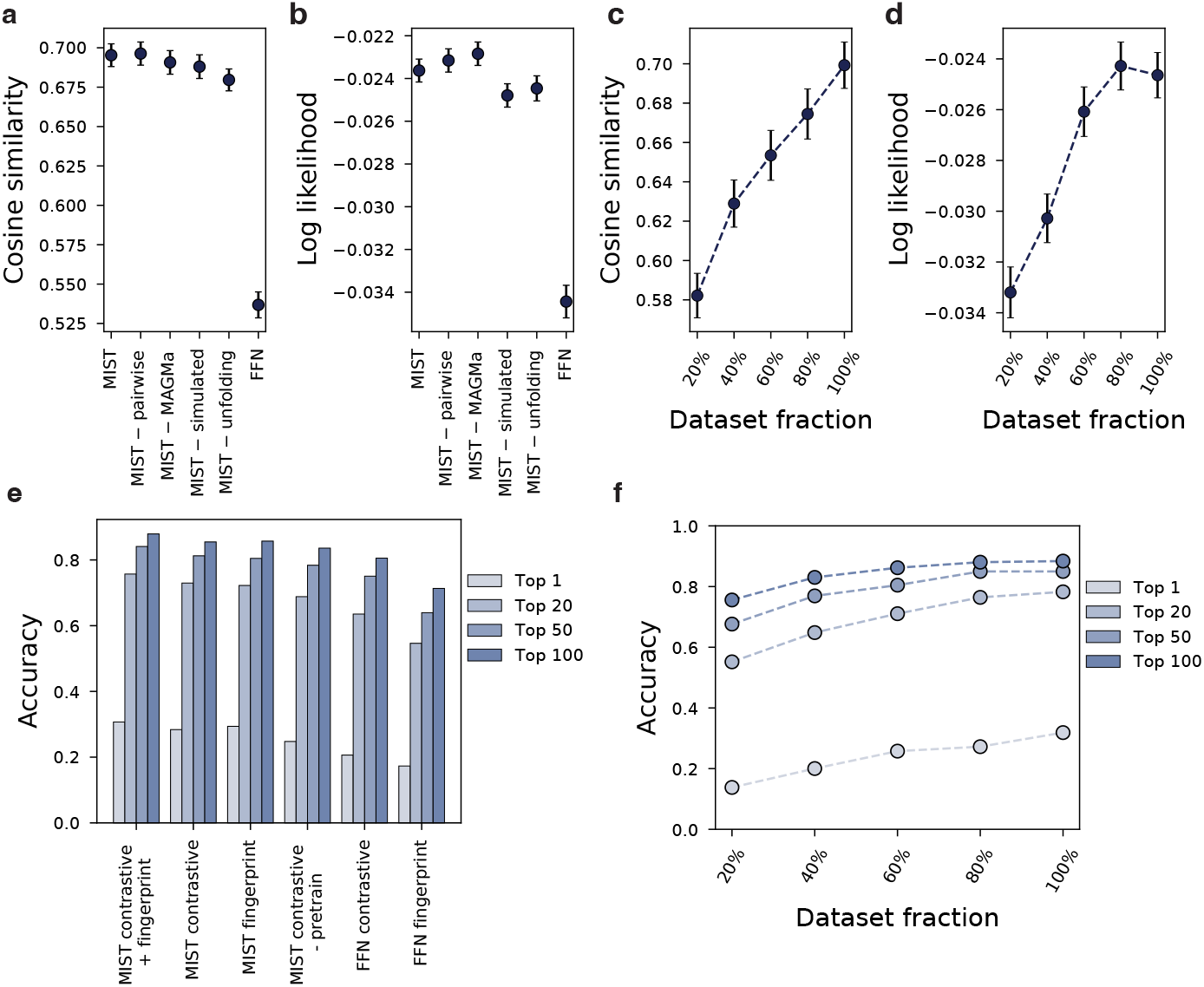
Model ablations affirm the value of domain-inspired model components. **a,b.** MIST performance on fingerprint prediction compared to ablated variants of MIST without pairwise interactions, without MAGMa substructure supervision, using no simulated spectra, without fingerprint unfolding and using only a feed forward network (FFN) using metrics of (**a**) cosine similarity and (**b**) log likelihood. Error bars show 95% confidence intervals for the mean. **c,d.** MIST fingerprint prediction accuracy improves as a function of dataset size for both (**c**) cosine similarity and (**d**) log likelihood. **e.** MIST retrieval performance using a weighted sum of contrastive and fingerprint distance outperforms contrastive distance or fingerprint distance alone, a contrastive version of the model without finetuning, a FFN contrastive model, or FFN fingerprints. **f.** Full MIST contrastive + fingerprint retrieval accuracy improves with training set size. All results are computed on the public GNPS subset data released with CANOPUS. Model ablations are conducted with 3 random splits of the data by molecular structure; dataset ablations are conducted for a single split of the data. Log likelihood values are clamped to a minimum of −5.

Ablations measuring retrieval accuracy further support the utility of a contrastive-fingerprint hybrid retrieval distance. Retrieval accuracies are computed for various “top-*k*” values using different retrieval scoring functions including the full MIST contrastive function (“MIST contrastive + fingerprint”), only contrastive distance (“MIST contrastive”), only fingerprint distance (“MIST fingerprint”), contrastive distance computed with a model that was not pretrained on fingerprint distance (“MIST contrastive - pre-train”), contrastive distances for a feed forward network (“FFN contrastive”) and fingerprints computed with a FFN model (“FFN fingerprint”). The full MIST retrieval using a weighted average of contrastive and fingerprint distances performs better than either distance alone at 30.7% Top 1 accuracy and 75.7% Top-20 accuracy (Fig. 6e, Tab. S4). Curiously, when using Morgan fingerprints, contrastive and fingerprint retrieval accuracies when computed independently are near equivalent unlike the proprietary dataset case. This reaffirms that the choice of fingerprint may add additional biases to retrieval. Excitingly, we observe that MIST architecture Top-1% accuracy improves steadily as we increase the amount of training data (Fig. 6f), indicating that MIST should scale well as additional labeled data becomes available, a key benefit of such neural network approaches.

## 3 Discussion

We present a new computational method for the annotation of novel metabolites in untargeted metabolomics, MIST. Unlike prior methods which either use handcrafted kernels or domain-agnostic feed forward neural architectures, MIST utilizes a novel, mass spectra-specific end-to-end neural network architecture to represent mass spectra as a set of chemical formulae (fragments/peaks) and intensities. The architecture borrows insights from years of statistical analysis of mass spectra such as the use of neutral loss fragments and molecular sub-structures as additional training signals while remaining sufficiently flexible to adapt to its training data. Because MIST is completely open source, it is straightforward to substitute any proprietary mass spectra training data or target fingerprints to train a new model.

We demonstrate how MIST can be used both for predicting molecular structure fingerprints and also for learning a meaningful continuous representation, or latent space, with contrastive learning. For the task of fingerprint prediction, MIST achieves superior accuracy to prior methods. For the tasks of structural elucidation as retrieval and for organizing spectra into molecular networks, MIST’s learned contrastive embeddings prove exceptionally effective. Both tasks are useful in the prospective annotation of clinical microbiome data, as demonstrated through the example of annotation of putative and differentially abundant dipeptide molecules and alkaloid compounds.

A key difficulty in mass spectrometry model development is the lack of well standardized benchmarks, which have been essential to progress in machine learning tasks. CASMI competitions simultaneously evaluate chemical formula annotation alongside retrieval annotation, which is also a function of the database used for retrieval, making such comparisons difficult to fully deconvolute; by design, the relevant training data is not restricted or provided. To facilitate future progress in this field, we provide fully benchmarked model ablations on a small and tractable subset of public GNPS data for both annotation and fingerprint prediction. Such standards will enable better comparisons between models across studies.

A limitation of this work is that MIST is highly dependent on proper chemical formula assignments from MS1, an upstream problem that we rely on the SIRIUS platform to solve. Higher accuracy formula annotation and new methods for this task will be synergistic with MIST. MIST is currently only trained on annotated spectra, not unannotated spectra. Future work will explore how strategies such as pretraining can further improve the quality of structure elucidation models.

MIST provides a competitive neural solution to transform a mass spectrum and predicted chemical formula to a molecular fingerprint or latent space embedding for structural elucidation. Just as protein structure prediction has become powered almost completely by neural network models, we anticipate a similar shift in small molecule structure elucidation pipelines.

## 4 Acknowledgments

The authors thank J. Bradshaw, R. Mercado, R. Barzilay, M. Wang, J.C. Hütter, J. Pacheco, C. Tzouanas, M. Zhu, and D. Hitchcock for valuable feedback and discussions on the work. The authors are especially grateful to K. Duhrkop and S. Böcker for providing data to directly compare to their CSI:FingerID model and help utilizing the SIRIUS software. S.G. was funded by the Takeda Healthcare AI Fellowship and the Machine Learning for Pharmaceutical Discovery and Synthesis consortium.

## 5 Code availability

All code to replicate experiments, train new models, and load pre-trained models is available at https://github.com/samgoldman97/mist. The exact repository version used in this work has been archived with Zenodo [39].

## 6 Data availability

Public data used for benchmarking MIST models as processed by Duhrkop et al. [23] can be downloaded alongside our code with full directions included at https://github.com/samgoldman97/mist and ref. [39]. Data for NIST and head-to-head CSI comparisons is unavailable due to strict licensing rules around the NIST20 [47] dataset. Data to to repeat the retrospective study and reanalysis of IBD data can be retrieved from the MassIVE database at accessions MSV000084908 (raw data) and MSV000086509 (cohort info).

## 7 Author contributions

S.G. wrote the software and conducted experiments. G.H. adapted MAGMa sub-structure labeling for auxiliary model training. S.G., J.W., and C.W.C. conceptualized the project and designed model components. M.S. and R.X. provided prospective clinical data analysis support. S.G. and C.W.C wrote the manuscript. C.W.C supervised the work.

## 9 Online methods

We outline the core structure of our model and training details below, along with key MIST parameters and datasets utilized.

### 9.1 MIST architecture

#### Chemical formula inputs

Each fragmentation spectrum, *x* is parsed from a corresponding ‘.ms’ file and contains a precursor mass *M*_0_(*x*) for the full compound and a list of *N_p_* mass/charge, intensity tuples: *x* = [(*m*_1_, *I*_1_), (*m*_2_, *I*_2_), … (*m_N_p__*, *I_N_p__*)]. MIST assumes that a user has first determined the chemical formula for the precursor compound *M*_0_(*x*) and removed adduct information (i.e., using SIRUIS, ZODIAC, or other tools [40, 69]). MIST then attempts to assign each *m_i_* fragment peak a chemical formula subset from the full mass, filtering out all masses for which this conversion is not possible. In this work, SIRIUS [40] is used to perform this step but may be replaced with other approaches. Following formula assignment by SIRIUS, peaks are sorted in order of decreasing intensity and only the top 60 most intense peaks are considered for chemical formula labeling.

#### Formulae embedder

Every chemical formula sub-peak is represented as a vector 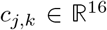, where each position in the vector is the integer count of atoms for common elements: *C, N, P, O, S, Si, I, H, Cl, F, Br, B, Se, Fe, Co*, and *As*, each normalized by the maximum observed atom count of that element in the dataset.

Each formula vector in the spectrum, including the root corresponding to *M*_0_(*x*), is embedded into a continuous fixed length vector with a shared shallow multi-layer perceptron (MLP) network:

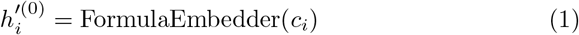

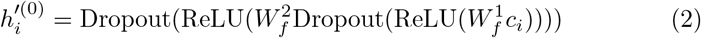

where 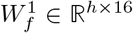 and 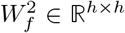 are learned parameters. Learned bias terms are used but omitted for clarity.

#### Set Transformer

Each embedded formula is concatenated with its relative intensity (normalized to be maximum 1 for each spectrum) as input to the transformer, 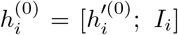, where [*a*; *b*] denotes a concatenation between vectors *a* and *b*.

A series of multi-headed attention layers [42] as in the transformer neural network architecture are applied to progressively update the hidden representation at each peak.

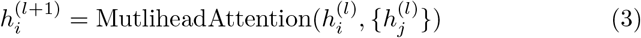

Unlike the original transformer, which uses positional encodings to maintain the ordering of inputs, we remove positional embeddings to make our model equivariant under permutations as in the SetTransformer[41]. Thus, each peak formula attends to other formulae in the spectrum to update its internal representation.

#### Pairwise attention

The relationship between each peak sub-formula in the spectrum can be explicitly written as the difference in atom counts between the two peaks. We incorporate this pairwise relationship into our model using featurized attention [70, 71].

Before entering the pairwise attention module, an all-by-all formula difference is computed for the input spectrum’s sub-formula list, *c* to extract these features. Formula differences are always positive and formula differences are only included when all differences have the same sign (i.e., one sub-formula is a superset of another):

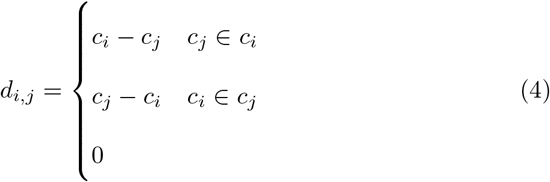

Each formula difference is embedded using a FormulaEmbedder module (shallow MLP) with parameters shared with the FormulaEmbedder used on the original peaks:

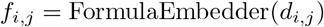

In the original transformer attention layers, attention is computed as

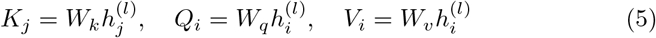

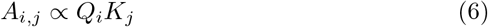

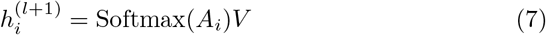

We incorporate attention features by modifying the attention scores to combine formula embeddings with formula difference embeddingas as peak and interaction representations, respectively:

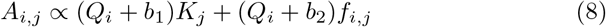

Here, *b*_1_ and *b*_2_ are trainable biases and *W_k_, W_q_, W_v_* are learned weight matrices.

#### Pooling

After iterating with *L* layers of multiheaded attention, a final hidden representation *h^out^* for the spectrum *x* is extracted using the hidden representation 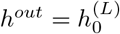, where the 0^*th*^ hidden state corresponds to the precursor mass formula. We empirically find that using the hidden representation at the precursor mass formula, rather than mean pooling over all peaks, greatly improves performance.

#### Loss function

The loss function to train MIST is composed of several different terms related to target fingerprint prediction 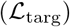, unfolding of fingerprints 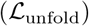, and substructure prediction 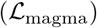. The relevant secondary terms are weighted by their corresponding hyperparameters λ_unfold_ and λ_magma_ to derive a loss for the full spectra:

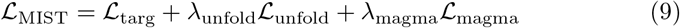

Each term is defined in depth below.

#### MAGMa substructure prediction

In addition to training MIST to predict full spectra fingerprints, we train MIST to predict 512-bit Morgan fingerprints for each labeled formula substructure as an additional training signal to regularize the model. For each training example composed of spectrum *x* and molecular structure *y*, we combinatorially enumerate possible substructures of *y* using the `MAGMa` algorithm to assign putative substructure fragments to peaks in *x* (Suplementary Information Fig. S4). We compute a Morgan fingerprint *q_j_* ∈ {0,1}^512^ for each of the *j^th^* sub-fragment [45]. We learn a separate prediction layer 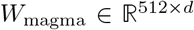 and apply this to the final representation *at each sub-peak* after the formula transformer layers:

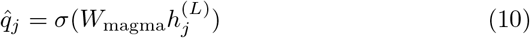

where *σ*(·) is the sigmoid activation function.

We derive an auxiliary substructure fingerprint loss from these predictions for all labeled sub-structures in each spectrum which empirically enhances performance:

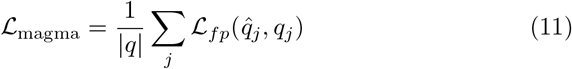

where 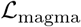 is a clipped cosine loss function, 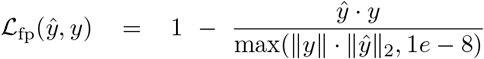. We note that the number of magma substructure labels is less than the number of peaks, since not all peaks have assigned substructures.

#### Fingerprint prediction by unfolding

The final representation 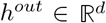 is used to predict the target binary fingerprint describing the molecular structure with dimension *D*: *y* ∈ {0,1}^*D*^. Previous approaches to fingerprint prediction have used simple feedforward networks from some representation of the spectrum, e.g., 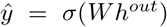. However, given the sparsity of our target fingerprint, we instead introduce an unfolding module to progressively grow the fingerprint by predicting *n*_grow_ folded fingerprint intermediates 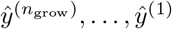, with a fingerprint loss calculation at each step. We deterministically define the folding function to establish targets, 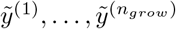, for each intermediate prediction layer, mirroring how hashed fingerprints like the Morgan fingerprint are folded into fixed-length vectors in the first place:

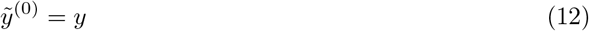

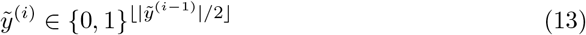

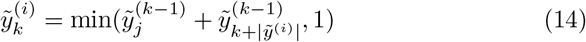

In doing so, each progressive target halves in length and bits are merged using the modulo function (i.e,. assuming there are *K* bits in the current fingerprint target, bit *k* is the 1 if either bit *k* or bit *k* + *K* is 1 in the previous vector). During model training, the network learns to invert this procedure such that the previous intermediate is first expanded then gated using learned weights 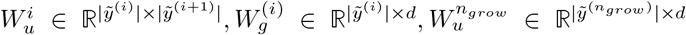. The lowest resolution target 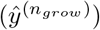 is predicted first and unfolded progressively:

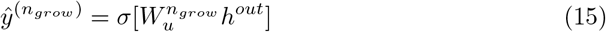

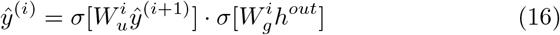

We define both a final fingerprint loss and unfolding loss corresponding to each layer:

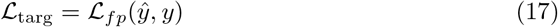

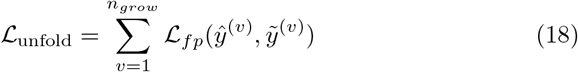

#### Forward model simulation

A key insight for MIST is to increase the total number of spectra in the dataset and, consequently, the diversity of fingerprints predicted. To do this, we train a “forward” simulator model to predict spectra from candidate molecule fingerprints similar to Wei et al. [31]. Using the same train and test split as MIST, we learn a model mapping from *y* to *x*, where *x* represents a spectrum and *y* is the same binary molecular fingerprint as in MIST. We convert *x* into a binned spectrum representation from m/z 0 to 1500 with bin spacings of 0.1 for a total of 15, 000 bins, *x* ∈ [0,1]^15000^. The intensity in each bin is normalized to [0,1] by dividing by the maximum intensity observed in that spectrum (Fig. S3a). Following Wei et al., we predict both fragment intensities and neutral losses and pool these together (Fig. S3d):

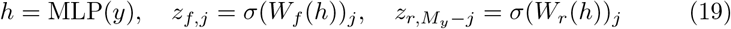

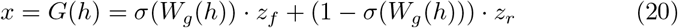

Here, MLP defines a shallow feed forward network transforming the input fingerprint into a hidden representation. *W_f_* and *W_r_* transform the hidden representation into predictions of peaks and losses. Losses are converted into a peaks by subtracting each predicted loss bin from *M_y_*, the bin of the full molecule mass. *W_g_* represents an affine trans formation from the hidden input into a set of gates to balance the forward and reverse prediction terms, *z_f_* and *z_r_*.

To further increase the performance of the forward simulator, we use a similar unfolding strategy as utilized in MIST for fingerprint prediction (Fig. S3c). We apply several repeated unfolding layers to predict *n_fg_* binned spectra with lower dimensions at each step 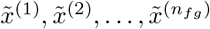. We define these by binning spectra targets with progressively lower resolution:

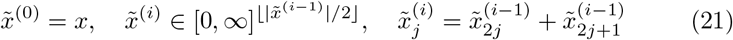

To make predictions, we first predict the lowest resolution 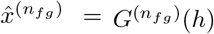. Following our strategy from fingerprint prediction, one strategy would be to predict the higher resolution spectrum 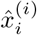 using a linear transformation of the coarser prediction 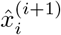. However, given the wide dimensionality, such learned transformations would have shape 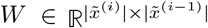. This is prohibitive with large bin sizes. Instead, we alternatively elect to naively expand 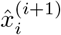 by setting 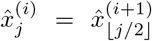 without any additional parametrization. We refine this prediction using refinement and weighting gates as a function of the original hidden representation 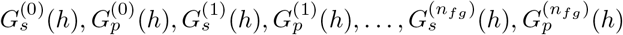:

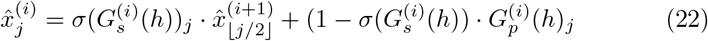

In doing so, each transformation 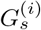 and 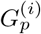 will involve weight matrices 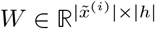, reducing the number of parameters substantially.

Spectra are filtered to include peaks at only the positions for which chemical sub-formula are labeled and sufficient intensities are observed. Unlike Wei et al, to train this model we use a binary cross entropy loss for the presence of certain fragments, rather than a cosine similarity loss to reduce the influence of low intensity peaks [31]. We define a binarized spectrum 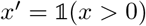 and also a weighting factor *s*(*x*) = Softmax(*x*) to enforce that peaks with higher intensities are weighted more heavily in our loss function:

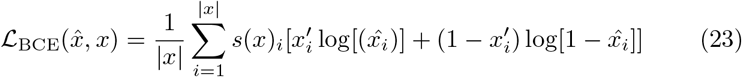

We apply this BCE loss at each unfolding layer step to get an unfolding loss and final target loss, weighted by a factor λ_forward_ to define the full forward loss:

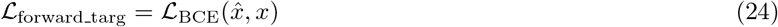

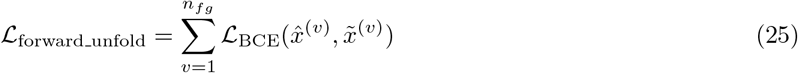

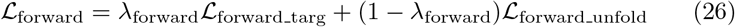

After training this model, we make spectra predictions for a randomly sampled set of biomolecules while excluding the validation and test set molecules to avoid accidentally biasing our model. We note that in additional experiments, including these molecules in the augmentation had a surprisingly strong effect in biasing our model toward the true predictions at test time, highlighting the ease for data leakage to inflate performance in this setting. For each predicted spectrum, we convert the binned spectrum back to sub-formula labels using combinatorial enumeration and using a maximum of 50 labeled sub-formulae with the highest predicted intensity. We use a prediction threshold of 0.2 to call peaks (Fig. S3b).

During training, in each epoch we include the entire training set of experimental spectra. We augment the training set in that batch to include a fraction of 1 – *f*_real_ simulated spectra, with *f*_real_ set during hyperparameter optimization, such that experimental spectra only make up a fraction *f*_real_ of each epoch.

#### Spectra noising

To make the model robust to peak shifts, each fragmentation spectrum is noised using a similar protocol to MetFID [36]. During training, with a *p*_noise_ = 0.5 probability any given spectrum is augmented. For each spectrum selected for augmentation, we define two augmentation operations: `remove`, in which a peak is fully removed, and `intensity`, in which a peak’s intensity is multiplied by a randomly sampled scalar value 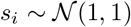, truncated to a minimum of 0. The probability of removing each peak, *p_j_* is proportional to its intensity value to ensure high intensity peaks are less likely to be removed:

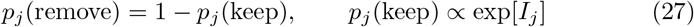

For each noised spectrum, we sample the number of peaks to remove from a binomial distribution with *p*_remove_ = 0.5 and we sample a second binomial distribution for the number of peaks to rescale with probability *p*_intensity_ = 0.1. Full pseudocode is listed in the Supplementary Information (Section 10.3).

### 9.2 Fingerprints

To directly compare against CSI:FingerID, we train models using the custom length 5,496 fingerprints included with SIRIUS as provided by the authors and computed using the Docker image provided for MSNovelist [25]. For ablation datasets on public data, we use `RDKit` to compute 4096-bit circular Morgan fingerprints [68, 72].

### 9.3 Feed forward baseline

We train a feed forward network baseline similar in concept to MetFID that maps binned spectra to fingerprints [36]. We bin mass values for each spectrum into *n*_bins_ equally spaced bins from 0 to 1500. *n*_bins_ is set in hyperparameter optimization. The model is trained using cosine similarity loss between the predicted fingerprint and the true fingerprint as in MIST’s architecture.

### 9.4 Contrastive fine-tuning for database retrieval

After pre-training MIST to predict fingerprints, we extract the model weights used to produce the hidden representation *h^out^* = *F*(*x*) and treat this internal representation as a contrastive space to conduct retrieval, where *F* defines the pretrained MIST feature extractor. We learn a separate single layer projection 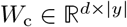 to map fingerprints into the contrastive space.

We sample a set of *n*_decoy_ decoy training fingerprints, *z*_1_, *z*_2_, … *z*_*n*_decoy__ for each spectrum and learn to project these into the latent space such that the true target fingerprint maps closer to the latent *h^out^* than to the decoys. We utilize a softmax contrastive loss inspired by noise contrastive estimation with temperature parameter *τ* and cosine similarity function [52, 73]:

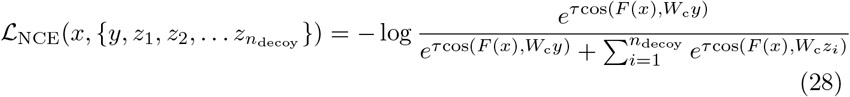

To sample decoys, we utilize the PubChem database (April 2022) to identify isomers with high similarity ot the ground truth for all spectra in the training set. We calculate the 256 closest examples for each isomer by Tanimoto similarity using Morgan fingerprints with length 4096 and radius 2. In each batch, these are sampled proportional to an exponential of their their Tanimoto similarity to the target:

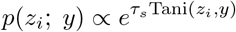

Similarity is weighted by an additional temperature factor *τ_s_* =4 enforcing that decoys with high Tanimoto similarity are sampled more often.

We include forward simulated spectra in contrastive fine-tuning as well. Rather than sample isomeric decoys from PubChem for these simulated examples, we extract decoy fingerprints from the set of all forward simulated molecules to avoid further additional computational complexity. We again sample decoys using Tanimoto similarity for hard negative mining.

During contrastive learning we train the network using a weighted sum of the contrastive loss and MIST loss with a hyperparameter λ_*c*_:

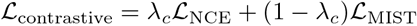

### 9.5 Model training

All models are trained with the RAdam [74] optimizer. Learning rate and weight decay hyperparameters are set for each dataset and detailed in Section 9.9. A validation set containing 5% of the data is excluded from the training set and utilized for early stopping during training with a patience of 20.

### 9.6 Retrieval

#### Fingerprint retrieval

Retrieval results are calculated using the PubChem database (April 2022) unless otherwise stated. For each spectrum, a fingerprint is first predicted. All isomers matching the spectrum’s chemical formula are rank ordered by cosine similarity between the isomer fingerprint and the predicted fingerprint. For all ties, the optimistic lower rank of the tied options is chosen. Ties are broken by selecting the minimum rank.

#### Bayesian retrieval

CSI:FingerID and SIRIUS internally utilize an adjusted distance, rather than cosine similarity, that corrects for correlations between fingerprint bits [51]. We utilize rankings calculated with Bayes as provided for CSI:FingerID predictions by the authors for direct comparison.

#### Contrastive retrieval

Contrastive retrieval results are computed by projecting fingerprints and spectra into the same continuous latent space using the trained contrastive model. We use a cosine distance metric in the latent space to rank isomers as in fingerprint retrieval. We ensemble contrastive distance and fingerprint distances with hyperparameter λ_*r*_ = 0.3, controlling the relative weights of fingerprint distance *d_fp_*(*x,y*) and contrastive distance 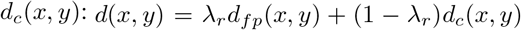. This parameter was manually tuned on a single data fold.

### 9.7 Training datasets

#### CSI:FingerID NIST comparison

We directly compare against CSI:FingerID using an [M+H]+ adduct dataset compiled from public data sources including the GNPS, MONA, and NIST20 [49]. This dataset has a total of 31, 145 spectra and 27, 797 unique compounds. We use cross validation, training on 3 independent structure-disjoint splits of the data.

#### GNPS CANOPUS benchmark

We utilize a public subset of the data including spectra pulled from the GNPS and MONA. This dataset is further filtered to [M+H]+ spectra and contains a total of 8,030 spectra and 7,131 unique compounds. We create structure-disjoint splits of this dataset using varying fractions of the dataset in the training set, 0.2, 0.4, 0.6, 0.8. For each of these settings, 5% of the training set is dynamically set aside for validation to perform early model stopping. This dataset is freely available and provided with the CANOPUS software tool [23].

#### Biomolecules

For forward simulation, we use biomolecules extracted from KEGG, KNAPSAcK, HMDB, and others [75, 76, 77], again prepared by Duhrkop *et al*. [23]. This dataset is only provided with InChIKey values, which cannot be mapped to molecular structures in the absence of a database lookup. We query PubChem to assign structural identities for a total of 997, 554 unique structures. We randomly sample this dataset to include 300,000 molecules (10x the full training dataset). Further, in each split, we exclude any molecules also appearing in the test and validation sets.

#### PubChem

PubChem was downloaded in its entirety [78] in April 2022 in order to build retrieval HDF5 files for chemical isomers. Comparisons to CSI:FingerID using PubChem as a retrieval library were made using a version of the PubChem database provided by the CSI:FingerID authors at the same time.

### 9.8 Clinical data reanalysis

Stool metabolite samples were previously collected by Mills et al. [56]. Metabolomic data was accessed under MassIVE accession number MSV000084908, including patient metadata at MSV000086509. Mass spectrometry data was extracted and converted into relative abundance tables and MS2 metabolite files using MZMine3 [79] to be used by SIRIUS (chemical formulae identification) and MIST (structure elucidation). MZMine3 parameters were set as specified by Mills et al., and an MS2 ppm tolerance of 10 was used for SIRIUS to identify chemical formulae.

MIST annotations were made using ensembles of 5 models. Structural annotations were first retrieved from the HMDB compound library; for all spectra without isomers in HMDB (i.e., no molecules in HMDB share the putative chemical formula), PubChem was utilized as a compound library for annotations.

Correlation coefficients with disease severity were computed within UC and CD cohorts for all patients that were assigned disease activity levels. Healthy-UC and healthy-CD comparisons were computed using independent two sided t-tests and adjusted with the Benjamini-Hochberg method. To utilize more equal sized cohort comparisons, UC and CD were subsetted to include patients with disease severity > 0.2 down to 40 and 22 patients respectively when comparing to healthy control groups.

### 9.9 Hyperparameter optimization

The tree-structured Parzen estimator hyperparameter search algorithm [80] as implemented by Optuna [81] was used to identify the best set of hyperparameters for each model. Hyperparameters were optimized once to maximize performance on a single validation split of the data, excluding auxilary loss terms. RayTune [82] was used to enable efficient parallelization across 3 RTX 3090 GPUs and a HyperBand scheduler was used to prune unpromising hyperparameters for efficiency. We ensured the same compute resources were available for tuning our models and each of the baselines and therefore capped resources at 100 trials on 3 GPUs. We conducted two separate hyperparameter sweeps to identify model parameters best for predicting fingerprints from CSI:FingerID (NIST dataset) and also the Morgan 4096 bit fingerprints (CANOPUS benchmark dataset). Full hyperparameter sweep parameters are listed in the Supplementary Information Tables S5 and S6.

### 9.10 Chemical classifications

NPClassifier [55] was used to produce chemical classifications of each compound. We query the public and easily accessed GNPS web server endpoint [49] and extract the superclass definition for chemical classification. Compounds with no predicted annotation are labeled as “Unknown.”

### 9.11 Latent distance comparisons

Spectra are projected into a shared spectrum-structure latent space using the fine-tuned contrastive MIST model. To assess the extent to which spectral similarity accurately reflects structural (Tanimoto) similarity, we repeat analyses from Spec2Vec [28] within a single fold of test data. We calculate all pairwise spectral similarities using cosine similarity and sort from highest to lowest similarity. At each threshold, we compute the average Tanimoto structural similarity. We compare to an arbitrary theoretical upper bound if paired spectra were sorted by Tanimoto similarity directly.

#### Spec2Vec

The pretrained Spec2Vec model is downloaded and directly used to embed all spectra into latent space. We repeat the same analysis above using Cosine similarity in Spec2Vec latent space.

#### Modified cosine similarity

Modified cosine similarity [12] is computed between spectra themselves using the `MatchMS` Python library [54].

#### UMAP computation

UMAPs for projected compounds are computed using the UMAP Python package applied to latent contrastive spectra embeddings in the test set [83].

## 10 Supplementary information

### 10.1 Additional model details

#### Unfolding layers

Novel unfolding modules are introduced in MIST to progressively grow the fingerprint prediction. Target fingerprints are compressed using modulo operations to derive “lower resolution” targets. These lower dimensional vectors inherently have collisions for properties. That is, rather than each bit indicating the presence of a single property or motif, each bit represents the presence of one of several properties (Fig. S1).

**Fig. S1.**
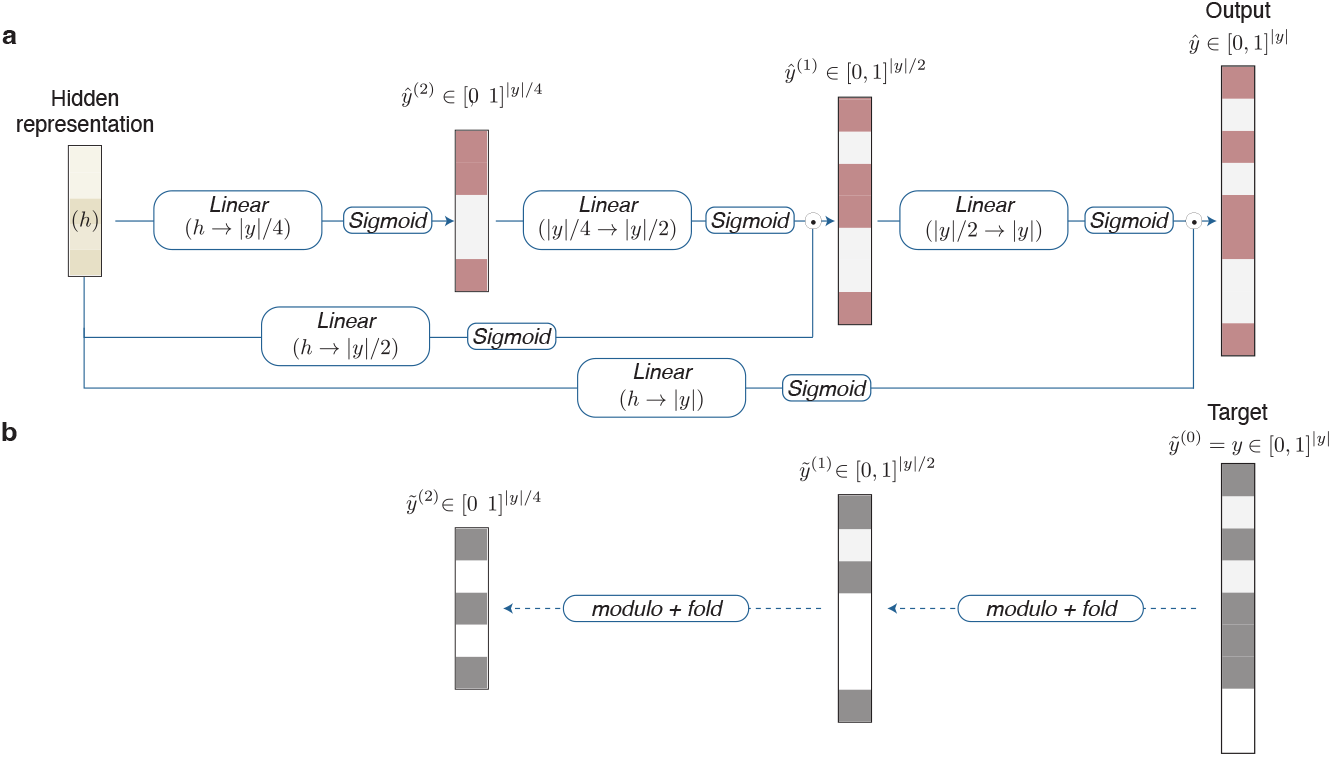
Unfolding fingerprint prediction module. **a.** The module starts with a hidden representation and subsequently predicts increasing resolutions of the fingerprint. **b.** At each intermediate fingerprint prediction 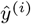, a corresponding target vector is generated by “folding” the previous, higher resolution fingerprint. An intermediate loss is calculated at each of these resolutions.

#### Binned feed forward baselines

A simple feed forward network inspired by MetFID is utilized as a baseline (Fig. S2) [36].

**Fig. S2.**
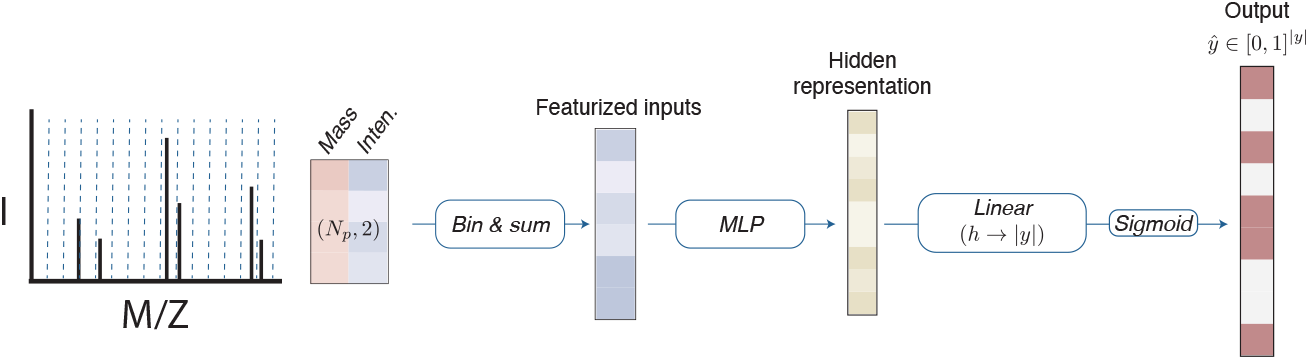
Feed forward network binned spectra baseline. An input spectrum is binned, a multilayer perceptron (MLP) is applied, and the output is projected to a fingerprint prediction.

### 10.2 MAGMa substructure labels

The MAGMa algorithm is used to take pairs of (molecule, spectrum) from the trained dataset and attribute molecule substructures to MS2 peaks [45]. In brief, the algorithm works by progressively removing atoms from the original structure to create substructures. Each time an atom is removed, the resulting subfragments are given a tolerance of ±*H* to account for hydrogen rearrangements. We highlight several spectra for the public GNPS dataset (Fig. S4).

**Fig. S3.**
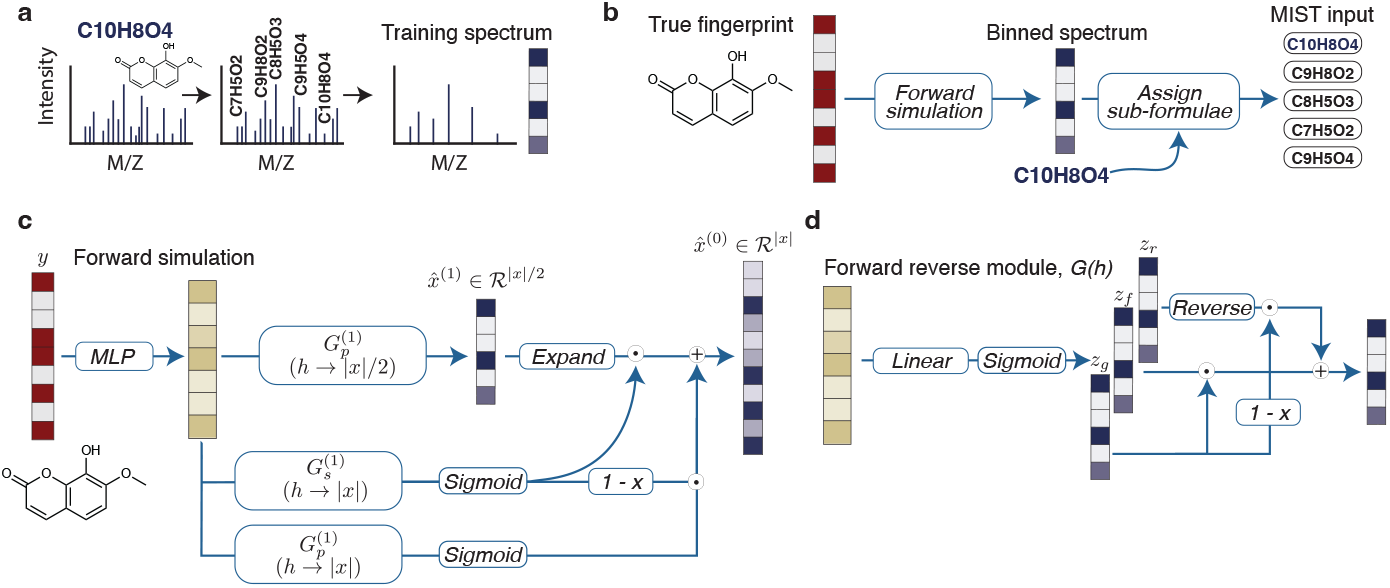
Forward simulation of spectra from molecules. **a.** An input spectrum, molecule pair is first denoised and cleaned by labeling chemical sub-formulae. This is converted into a binned spectra representation. **b.** To generate new model training examples, molecules are fingerprinted, binned spectra predictions are predicted by a neural network, and sub-formula are assigned to each of the bins such that the spectra can be used as input to MIST. **c.** The forward simulation neural architecture utilizes unfolding layers. A multilayer perceptron (MLP) projects the molecular fingerprint into a hidden representational space. Forward-reverse models, *G*, are used to progressively grow the binned prediction such that resolution increases at each step, similar to unfolding layers. **d.** Illustration of the forward reverse module, where intensities (*z_f_*) and neutral losses (*z_r_*) are predicted. Neutral losses are reversed and mapped onto intensity bins using the full mass as input then summed according to a learned gate (*z_g_*) to get a single prediction of intensities in each bin.

### 10.3 Spectra noising

To increase generality as in MetFID [36], we introduce an algorithm to probabilistically add noise to input training spectra (Alg. S1).

**Fig. S4.**
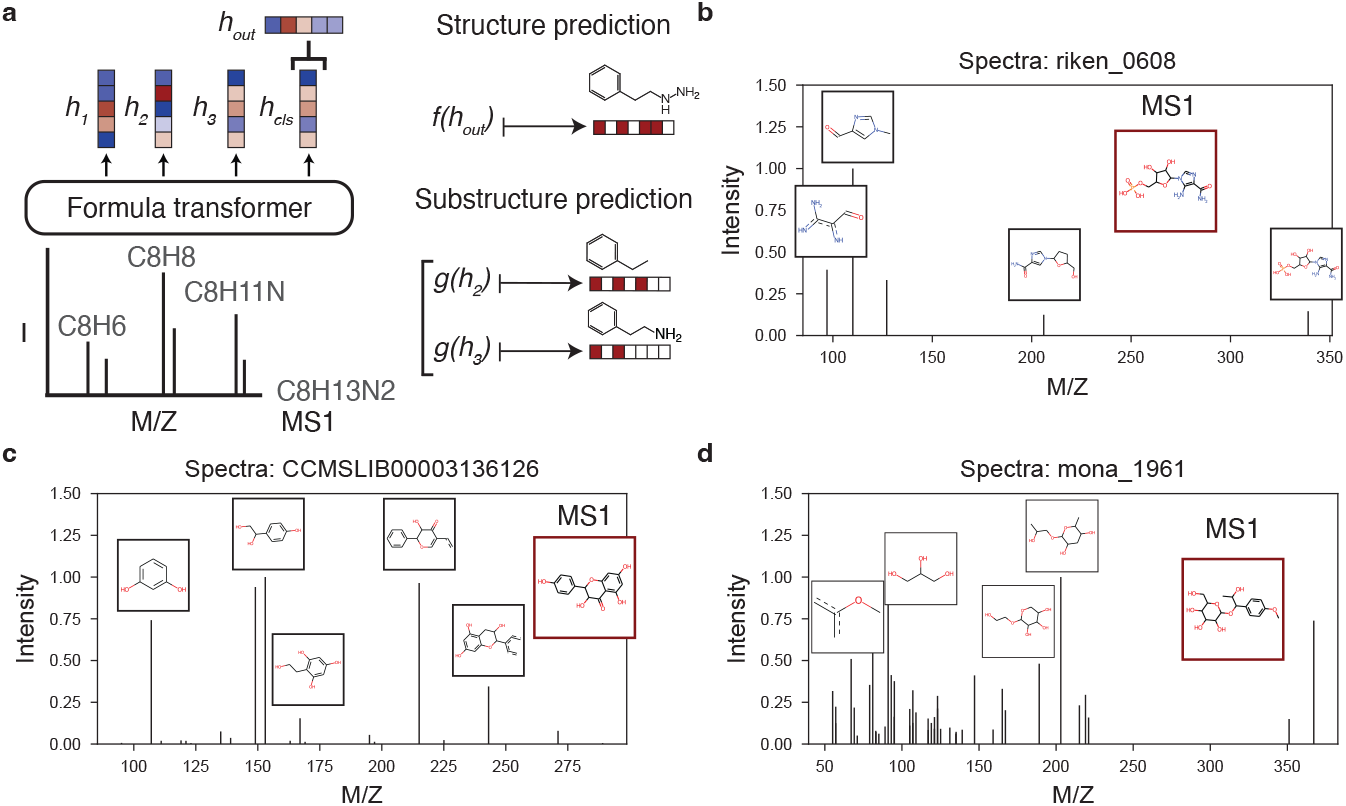
Labeling data substructures. **a.** MIST is trained to build representations of each peak in the spectrum. The model uses the MS1 peak to predict a fingerprint of the full molecule, and when substructures can be labeled for other peaks, the model learns to predict these as well. **b-d.** Example substructure peak labels in the training set computed with MAGMa. The full compound is outlined in red. Other substructures for the top 6 highest intensity peaks are outlined in black.

**Table S1.**
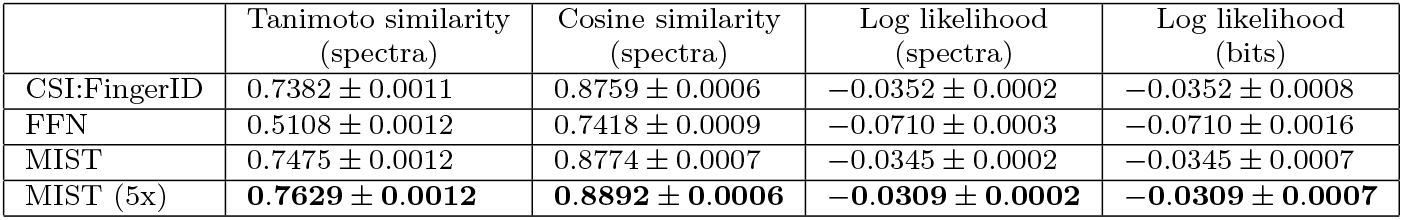
Full performance metrics for CSI:FingerID, FFN, MIST, and a MIST ensemble of 5 models are shown averaged across 3 structural splits of the data. Log likelihoods are clamped such that they have a minimum of −5. Results are shown ± standard error with an arbitrary number of significant figures.

**Table S2.** Full retrieval accuracy performance metrics for CSI:FingerID, FFN, MIST, and a MIST ensemble of 5 models are shown averaged across 3 structural splits of the data.

|  | Top 1(%) | Top 5(%) | Top 10(%) | Top 20(%) | Top 50(%) | Top 100(%) | Top 200(%) |
| --- | --- | --- | --- | --- | --- | --- | --- |
| CSI:FingerID + Cosine | 38.24 | 68.88 | 77.07 | 83.20 | 89.12 | 92.23 | 94.76 |
| CSI:FingerID + Bayes | **43.00** | **74.77** | **82.15** | 87.23 | 91.70 | 94.13 | 96.00 |
| MIST + Cosine | 29.72 | 59.56 | 69.26 | 73.60 | 84.10 | 88.13 | 91.84 |
| MIST + Contrastive | 34.45 | 69.28 | 79.04 | 85.75 | 91.63 | 94.56 | 96.49 |
| MIST (5x) + Cosine | 31.70 | 62.85 | 72.33 | 79.05 | 85.92 | 89.76 | 93.00 |
| MIST (5x) + Contrastive | 37.39 | 72.90 | 82.03 | **88.20** | **93.32** | **95.75** | **97.40** |

#### Algorithm S1 Noising a training spectrum

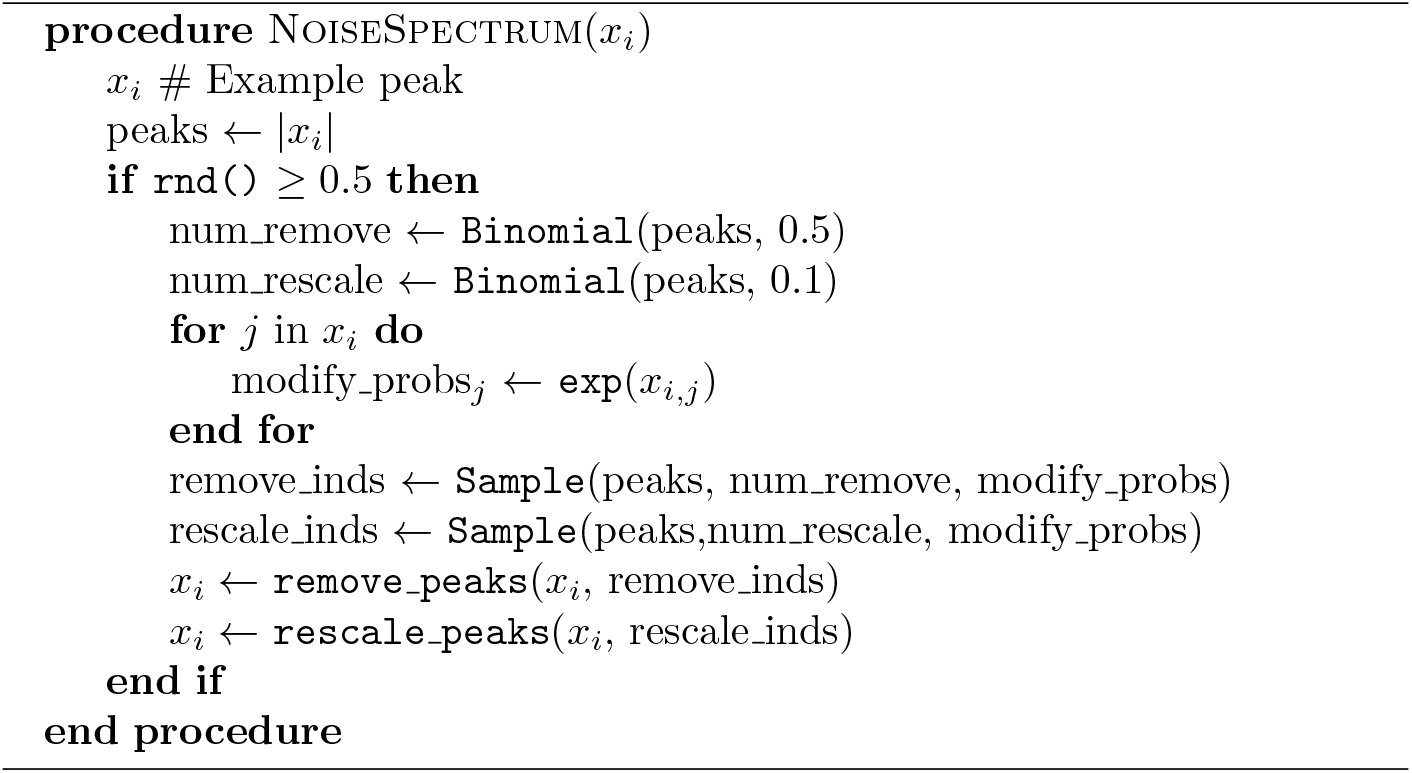

**Fig. S5.**
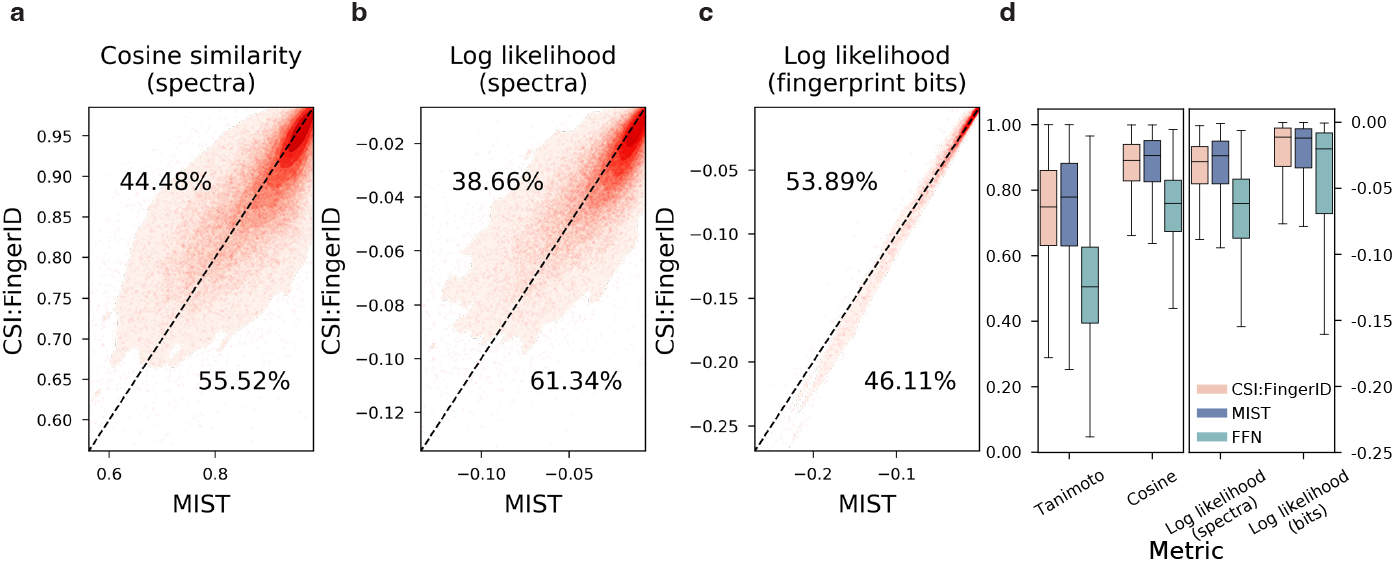
Single MIST model comparison to CSI:FingerID. Molecular fingerprints are predicted by MIST and CSI:FingerID for every spectrum in the test set using a single MIST model as in Figure 2. The performance for each spectrum by cosine similarity to the true fingerprint **(a)** or log likelihood **(b)** is evaluated and plotted. Points below the line represent instances where MIST is more performant. **c.** Equivalent evaluation showing the likelihood of predicting each fingerprint bit correctly across all spectra. **d.** The performance of CSI:FingerID, MIST, and FFN, a baseline inspired by MetFID, are shown. “Tanimoto,” “Cosine,” and “Log likelihood (spectra)” indicate performance across spectra and “Log likelihood (bits)” indicates performance across bits (higher is better). Median lines are shown; boxes show the interquartile range, with fliers indicating 1.5x interquartile ranges. Results are aggregated across 3 splits of the data.

**Table S3.** Full fingerprint accuracy accuracy on CANOPUS GNPS subset dataset for various model ablations. All model performances are computed on a merged dataset of fingerprint predictions computed for 3 independent random structural splits of 3 random structural splits of the CANOPUS dataset. FFN: Feed forward neural network.

| Model | Cosine similarity | Log likelihood |
| --- | --- | --- |
| FFN | 0.5368 | -0.0344 |
| MIST - unfolding | 0.6796 | -0.0244 |
| MIST - simulated | 0.6880 | -0.0248 |
| MIST - MAGMa | 0.6907 | **-0.0228** |
| MIST - pairwise | **0.6963** | -0.0232 |
| MIST | 0.6953 | -0.0236 |

**Table S4.** Full retrieval accuracy on CANOPUS GNPS subset dataset for various model ablations. All model performances are computed on a merged dataset of retrieval predictions computed for 3 independent random structural splits of 3 random structural splits of the CANOPUS dataset. FFN: Feed forward neural network.

|  | Top 1(%) | Top 5(%) | Top 10(%) | Top 20(%) | Top 50(%) | Top 100(%) | Top 200(%) |
| --- | --- | --- | --- | --- | --- | --- | --- |
| FFN fingerprint | 17.309 | 37.121 | 45.611 | 54.634 | 63.946 | 71.329 | 77.359 |
| FFN contrastive | 20.632 | 44.996 | 54.389 | 63.536 | 75.062 | 80.558 | 86.095 |
| MIST contrastive - pretrain | 24.774 | 50.656 | 60.624 | 68.827 | 78.384 | 83.593 | 88.146 |
| MIST contrastive | 28.384 | 55.373 | 65.217 | 72.970 | 81.255 | 85.480 | 89.377 |
| MIST fingerprint | 29.368 | 55.332 | 63.536 | 72.231 | 80.476 | 85.726 | 89.418 |
| MIST contrastive + fingerprint | **30.703** | **58.120** | **68.927** | **75.709** | **84.094** | **87.916** | **92.355** |

**Table S5.**
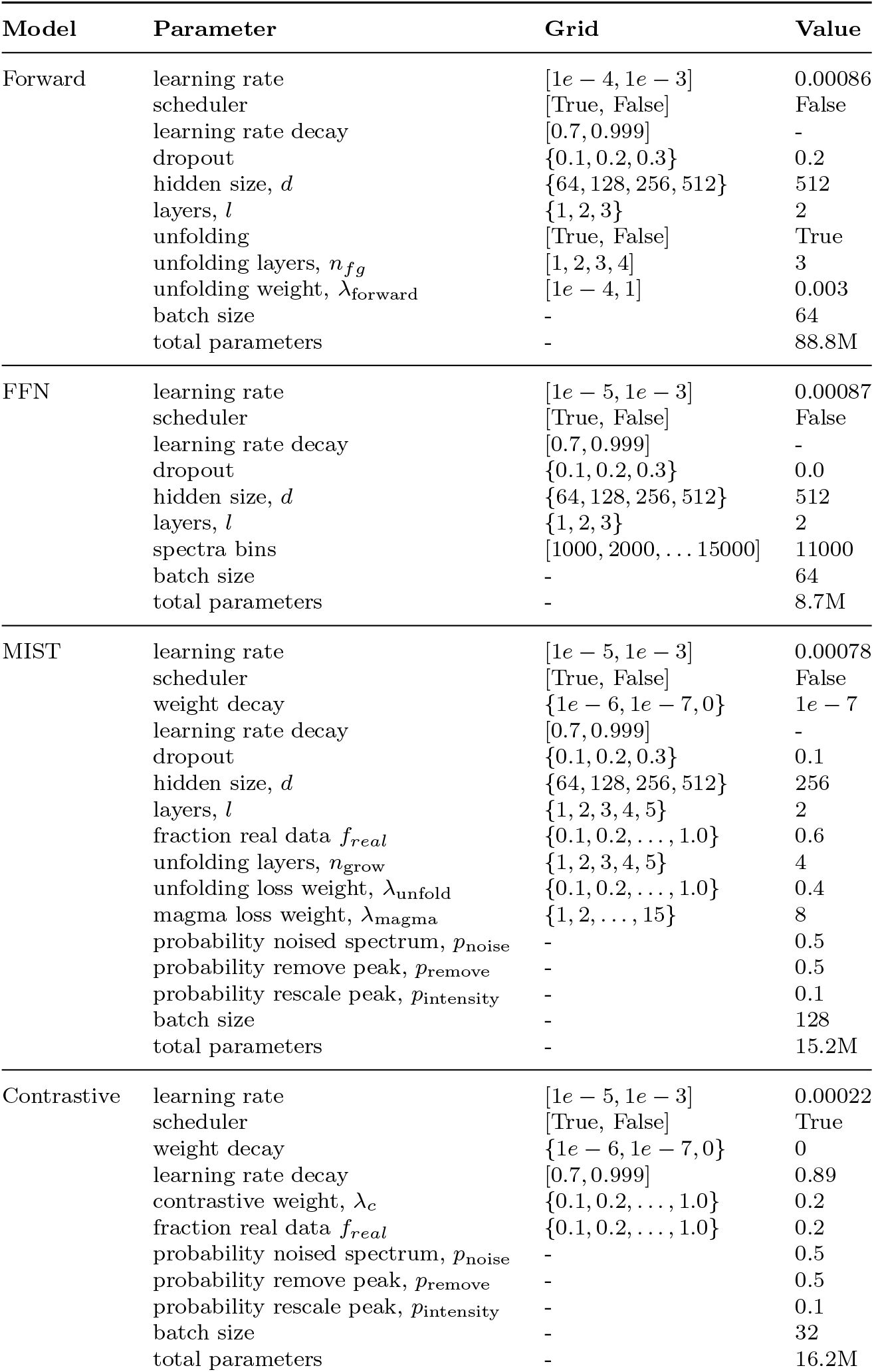
Model hyperparameters searched and set for CSI fingerprint prediction.

**Table S6.** Model hyperparameters searched and set for Morgan fingerprint prediction.

| Model | Parameter | Grid | Value |
| --- | --- | --- | --- |
| Forward | learning rate | $[1e-4, 1e-3]$ | 0.00054 |
|  | scheduler | [True, False] | False |
|  | learning rate decay | [0.7, 0.999] | - |
|  | dropout | {0.1, 0.2, 0.3} | 0.3 |
| | hidden size, $d$ | {64, 128, 256, 512} | 512 |
| | layers, $l$ | {1, 2, 3} | 2 |
|  | unfolding | [True, False] | True |
| | unfolding layers, $n_{fg}$ | [1, 2, 3, 4] | 3 |
| | unfolding weight, $\lambda_{\text{forward}}$ | $[1e-4, 1]$ | 0.0016 |
|  | batch size | - | 64 |
|  | total parameters | - | 83.7M |
| FFN | learning rate | $[1e-5, 1e-3]$ | 0.00087 |
|  | scheduler | [True, False] | False |
|  | learning rate decay | [0.7, 0.999] | - |
|  | dropout | {0.1, 0.2, 0.3} | 0.3 |
| | hidden size, $d$ | {64, 128, 256, 512} | 512 |
| | layers, $l$ | {1, 2, 3} | 2 |
|  | spectra bins | [1000, 2000, ... 15000] | 11000 |
|  | batch size | - | 64 |
|  | total parameters | - | 8.0M |
| MIST | learning rate | $[1e-5, 1e-3]$ | 0.0003 |
|  | scheduler | [True, False] | False |
| | weight decay | $\{1e-6, 1e-7, 0\}$ | $1e-7$ |
|  | learning rate decay | [0.7, 0.999] | - |
|  | dropout | {0.1, 0.2, 0.3} | 0.0 |
| | hidden size, $d$ | {64, 128, 256, 512} | 512 |
| | layers, $l$ | {1, 2, 3, 4, 5} | 3 |
| | fraction real data $f_{\text{real}}$ | {0.1, 0.2, ..., 1.0} | 0.1 |
| | unfolding layers, $n_{\text{grow}}$ | {1, 2, 3, 4, 5} | 5 |
| | unfolding loss weight, $\lambda_{\text{unfold}}$ | {0.1, 0.2, ..., 1.0} | 0.1 |
| | magma loss weight, $\lambda_{\text{magma}}$ | {1, 2, ..., 15} | 4 |
| | probability noised spectrum, $p_{\text{noise}}$ | - | 0.5 |
| | probability remove peak, $p_{\text{remove}}$ | - | 0.5 |
| | probability rescale peak, $p_{\text{intensity}}$ | - | 0.1 |
|  | batch size | - | 128 |
|  | total parameters | - | 25.6M |
| Contrastive | learning rate | $[1e-5, 1e-3]$ | 0.0002 |
|  | scheduler | [True, False] | True |
| | weight decay | $\{1e-6, 1e-7, 0\}$ | $1e-7$ |
|  | learning rate decay (10k steps) | [0.7, 0.999] | 0.89 |
| | contrastive weight, $\lambda_{\text{c}}$ | {0.1, 0.2, ..., 1.0} | 0.2 |
| | fraction real data $f_{\text{real}}$ | {0.1, 0.2, ..., 1.0} | 0.2 |
| | probability noised spectrum, $p_{\text{noise}}$ | - | 0.5 |
| | probability remove peak, $p_{\text{remove}}$ | - | 0.5 |
| | probability rescale peak, $p_{\text{intensity}}$ | - | 0.1 |
|  | batch size | - | 32 |
|  | total parameters | - | 27.7M |

